# De novo engineered guide RNA-directed transposition with TnpB-family proteins reveals features of naturally evolved systems

**DOI:** 10.1101/2025.08.27.672191

**Authors:** Richard D. Schargel, Laura Chacon Machado, Shravanika Kumaran, Jordan E. Thesier, Alba Guarné, Joseph E. Peters

## Abstract

Programmable DNA integration using CRISPR-associated transposons (CASTs) offers powerful capabilities for genome engineering. The single effector Cas12k CAST examples evolved from a fixed guide TnpB nuclease protein. Here, we engineer de novo RNA-guided transposition systems, where the single guide RNA effector components are repurposed nuclease-dead TnpB-family proteins. These compact systems mediate high-efficiency guide RNA-directed DNA insertion with preserved orientation control and target immunity, reduced off-site targeting, release of a host factor requirement, and can be paired with an exonuclease domain to mediate cut-and-paste transposition. In this engineered context, the TnpB derivatives show features not predicted from the original enzymes suggesting untapped avenues for improvement. In parallel, we show that mutations at the TniQ-TnsC interface in the Cas12k CAST system selectively attenuate off-site insertions while enhancing on-site activity. These results establish how Cas12 proteins and antecedent TnpB proteins can be engineered for high performance and specificity with guide RNA directed systems.

## Introduction

CRISPR-associated transposons (CASTs) are naturally evolved Tn7 family elements that coopted diverse CRISPR effectors to mediate programmable RNA-guided transposition of multi-kilobase DNA cargo.^1–3^ This mechanism of targeted DNA integration operates without requiring a host-directed homologous recombination process to occur at a programmed double-stranded break in the target DNA, making CASTs promising candidates as genome editing tools. Many CAST elements have been identified, each occurring by independent acquisition events of different CRISPR-Cas systems by diverse members of Tn7 family elements.^1,4,6,45^ The CAST elements are commonly referred to by the CRISPR-Cas system subtype that was coopted by the transposon. Systems confirmed in a heterologous host include type V-K^2^, I-F3^3,5^, I-D^6^, and two distinct lineages from type I-B (referred to as I-B1 and I-B2).^7^ The type I examples use multi-subunit CRISPR-Cas complexes (Cascades), whereas in the type V-K systems a singular Cas12k effector possesses all the functions for targeting^8^. Although all CAST families support programmable DNA integration, the extensive coding size of type I systems^3,5–7^ may pose challenges for gene editing, especially in eukaryotes where delivery constraints are more pronounced.^56^ Type V-K CASTs are more compact in coding size but have other limitations with off-site targeting and a host factor requirement.^2^

Type V-K CASTs utilize four transposon-encoded proteins and a host factor to catalyze replicative transposition.^2,10,11,41^ Cas12k, the CRISPR effector, recognizes a target site complementary to its gRNA.^2,9^ The host factor, S15, interacts with the RNA scaffold and allosterically stimulates the recruitment of TniQ and TnsC.^2,10,11^ TnsC, a AAA+ ATPase, acts as a central regulator, coordinating the recruitment and activation of the TnsB transposase complex only upon target site recognition.^9,12^ TnsB, the DDE-family transposase, executes DNA strand cleavage and integration steps by mediating breakage and joining during transposition.^13,14^ Despite the apparent advantages of the type V-K system, limitations also exist such as high levels of effector-independent off-site transposition (TnsB+TnsC+TniQ only transposition). The V-K systems also use replicative transposition, a process shown to require double-strand break repair in related systems where it has been carefully characterized.^42,43^ The replicative transposition process also results in cointegrate insertions by duplication of the entire element and incorporation of the plasmid backbone at the target site.^41^ A subsequent processing event involving host homologous recombination is required to convert this structure into a single insertion in the target DNA. Previous work demonstrated that fusing nAniI nicking homing endonucleases to TnsB, the transposase, enables the non-transferred strand to be cleaved allowing cut-and-paste transposition, presumably bypassing the replicative transposition process that leads to contigrates.^14^

The Cas12 effectors evolved from guide RNA-directed nucleases, the TnpB proteins.^17–20^ A promising engineering avenue would be directly adapting TnpB proteins as transposon targeting modules. TnpB family proteins can be significantly smaller (∼400 amino acids) than Cas12k yet retain the programmability and target specificity of Cas12 effectors (**Figure 1A, Figure S1**).^21,22^ TnpB-family proteins are guided by a structured gRNA which consists of a 3’ short target-complementary sequence and a conserved 5’ scaffold that folds into a defined secondary structure for effector binding.^21–24^ Unlike CRISPR-Cas systems, which rely on arrays processed into individual guide RNAs, there is a single gRNA found in the TnpB-family systems encoded within and adjacent to their associated protein reading frame. Bioinformatic analyses reveal that TnpB homologs are widespread across both prokaryotic and eukaryotic organisms, exhibiting extensive structural and functional diversity.^17,20^ Among these, eukaryotic TnpB homologs, referred to as Fanzors, have gained significant interest as genome-editing tools, owing to their compact size and natural compatibility with eukaryotic cellular environments.^25,26^ Notably, a newly identified subset of TnpB homologs, called TnpB-like nuclease-dead repressors (TldRs), functions as RNA-guided transcriptional repressors.^27^ Remarkably, these proteins are even smaller (∼300-350 amino acids) than prototypical TnpBs and lack nuclease activity due to deletions in their catalytic domains (**Figure S1**).

**Fig 1:**
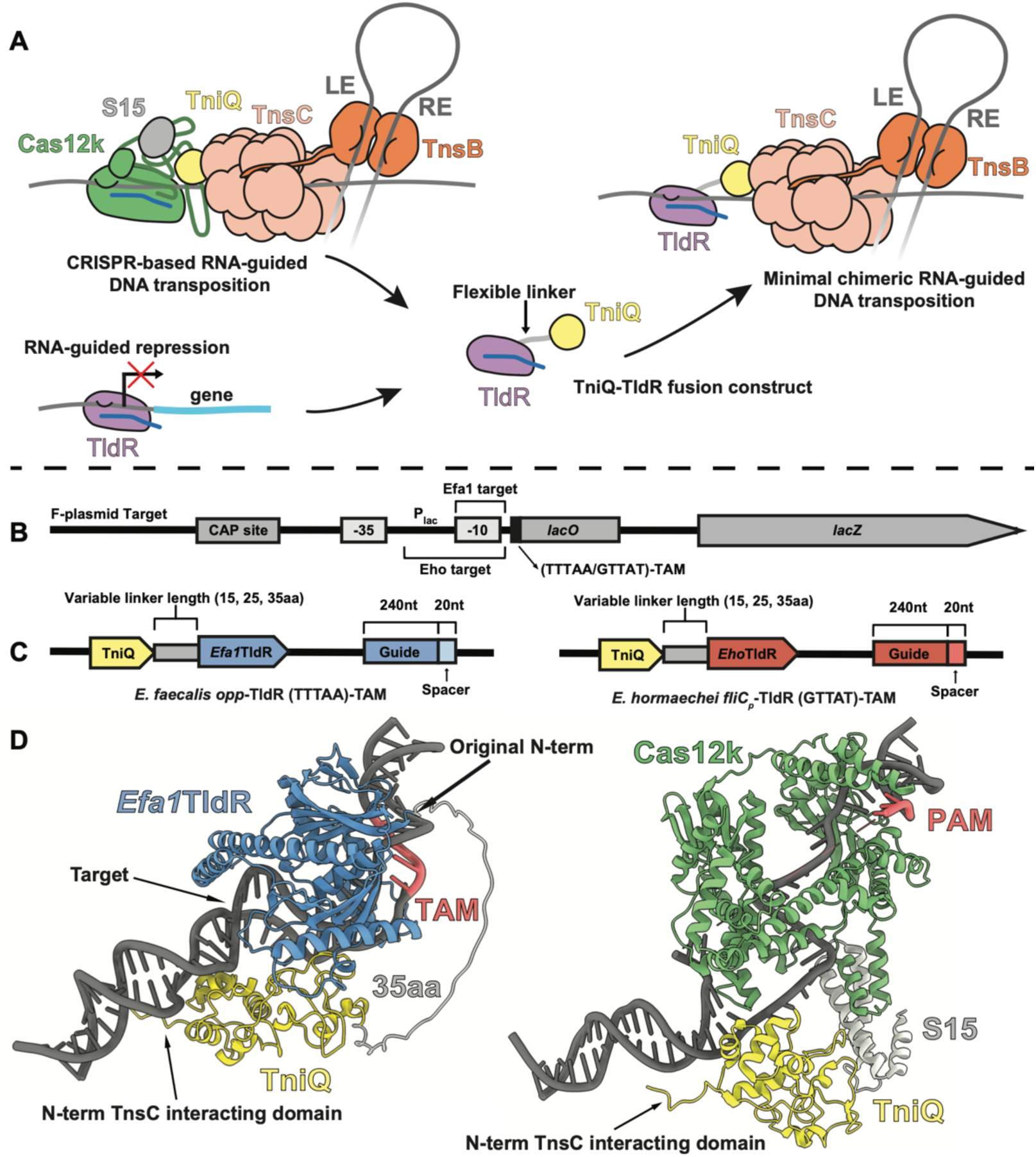
Development of *de novo* non-CRISPR RNA-guided transposition. **A.** Concept diagram of engineered TldR-guided transposition **B.** Schematic of TniQ-TldR fusion target construct. The lacZ gene was cloned into the F plasmid alongside its promoter and transcriptional regulation sites. Either TTTAA (*Efa1*TldR) or GTTAT (*Eho*TldR) TAM motifs were cloned directly upstream of the lacO site. **C**. Schematic of TniQ-Linker-Efa1 (left) and TniQ-Linker-Eho (right) construct. Guide sequence was designed to target P_lac_ sites shown in panel **B**. **D**. Structure of ShTniQ-35aa-Efa1 generated by AlphaFold3^28^ compared to ShCas12k-ShTniQ-S15 complex (8EA3)^10^ determined by cryoEM. ShQ-35aa-Efa1 prediction forms complex with DNA that is primed for recruitment of further transposition proteins. GuideRNA chains were removed for clarity.

Due to their minimal size, potential programmability, specificity, and relation to the Cas12 family of CRISPR effectors, we investigated representatives of the TnpB family (prototypical TnpB, Fanzors, and TldRs) as potential candidates for replacing Cas12k or other targeting modalities found with Tn7 family elements. We also address a limitation of the ShCAST V-K CAST elements with off-site effector-independent transposition revealing functional differences across V-K CAST relatives. We show that representatives from the two major branches of TldR can be engineered to produce de novo gRNA-directed transposition systems. This work provides a broader understanding of basic features of natural CAST systems and a greater understanding of the TnpB family that paves the way for more comprehensive use of these miniature programable DNA-binding proteins.

## Results

### ShTniQ-TldR fusions allow de novo RNA-guided transposition systems

Our initial work focused on engineering the transposon system associated with the V-K CAST family member from *Scytonema hofmannii* (ShCAST) based on its high activity, small size, and the considerable structural information available with this system. Our strategy involved replacing Cas12k with compact, programmable DNA-binding proteins by fusing ShCAST’s TniQ to members of the TnpB family. We focused on the TldR subgroup from the TnpB family because they naturally lack nuclease activity, functioning as guide RNA-directed DNA binding systems. Two major clades of TldRs have been characterized, each associated with repression of distinct host genes. The first branch, termed *fliC_p_*-TldRs are encoded in bacteriophages and contain gRNAs corresponding to *fliC* genes in *Enterobacteriaceae.*^27^ The second branch, termed *oppF*-TldRs, are encoded in *Enterococcal* genomes and possess gRNAs corresponding to multiple genes in the *opp* operon.^27^ Besides the native gRNA targets, these two branches possess different TAM (analogous to PAM in CRISPR-Cas systems) specificities, gRNA length requirements, and have different deletions when compared to canonical TnpB enzymes rendering them catalytically dead. For our initial round of TniQ-TldR fusions we selected a well-characterized representative from each branch; a *fliC_p_*-TldR from *Enterococcus faecalis* (*Efa1*TldR) and an *oppF*-TldR from *Enterobacter hormaechei* (*Eho*TldR) (**Figure 1, Figure S1**).

Structural predictions suggest that the N-terminus of TldR proteins is positioned distal to the downstream target DNA where transposition proteins would be recruited. As a result, a direct TniQ-TldR fusion without a flexible linker would likely restrict proper positioning of TniQ near the DNA target site. Thus, we designed our constructs to have linkers of various lengths (15, 25, and 35 amino acids) between TniQ and the TldR. Finally, we designed our initial gRNAs to target a TAM adjacent to the *lac* promoter (P_lac_) upstream of *lacZ* (**Figure 1B**). As an estimation of our constructs, we were able to generate AlphaFold3 predictions in which the linkers provided enough flexibility between the two proteins to position TniQ in the correct orientation to bind DNA and recruit subsequent transposition proteins (**Figure 1D**).^28^

For our initial experiment, we investigated whether our fusion constructs could be gRNA programmed to bind DNA to check if the ShTniQ fusion perturbed TldR targeting ability. To test this, we targeted P*_lac_* and measured whether our fusion constructs could repress β-galactosidase (β-gal) expression. Relative to our expression controls, we found that both the *Efa*1TldR and *Eho*TldR fusions substantially repressed β-gal activity (**Figure 2A**). For the ShTniQ-*Efa*1 fusions, we observed approximately 75% repression of β-gal activity across the three linker lengths. The ShTniQ-*Eho* fusions showed weaker and more variable repression ranging from approximately 60% to 45% for the 15aa and 25/35aa linkers, respectively. We also observed stronger repression with our ShTniQ-*Efa1* fusions relative to the WT *Efa1*TldR, with the inverse observed for the ShTniQ-*Eho* fusions. This result, however, may be related to varying levels of protein expression with ShTniQ fused to its N-terminus.

**Fig 2:**
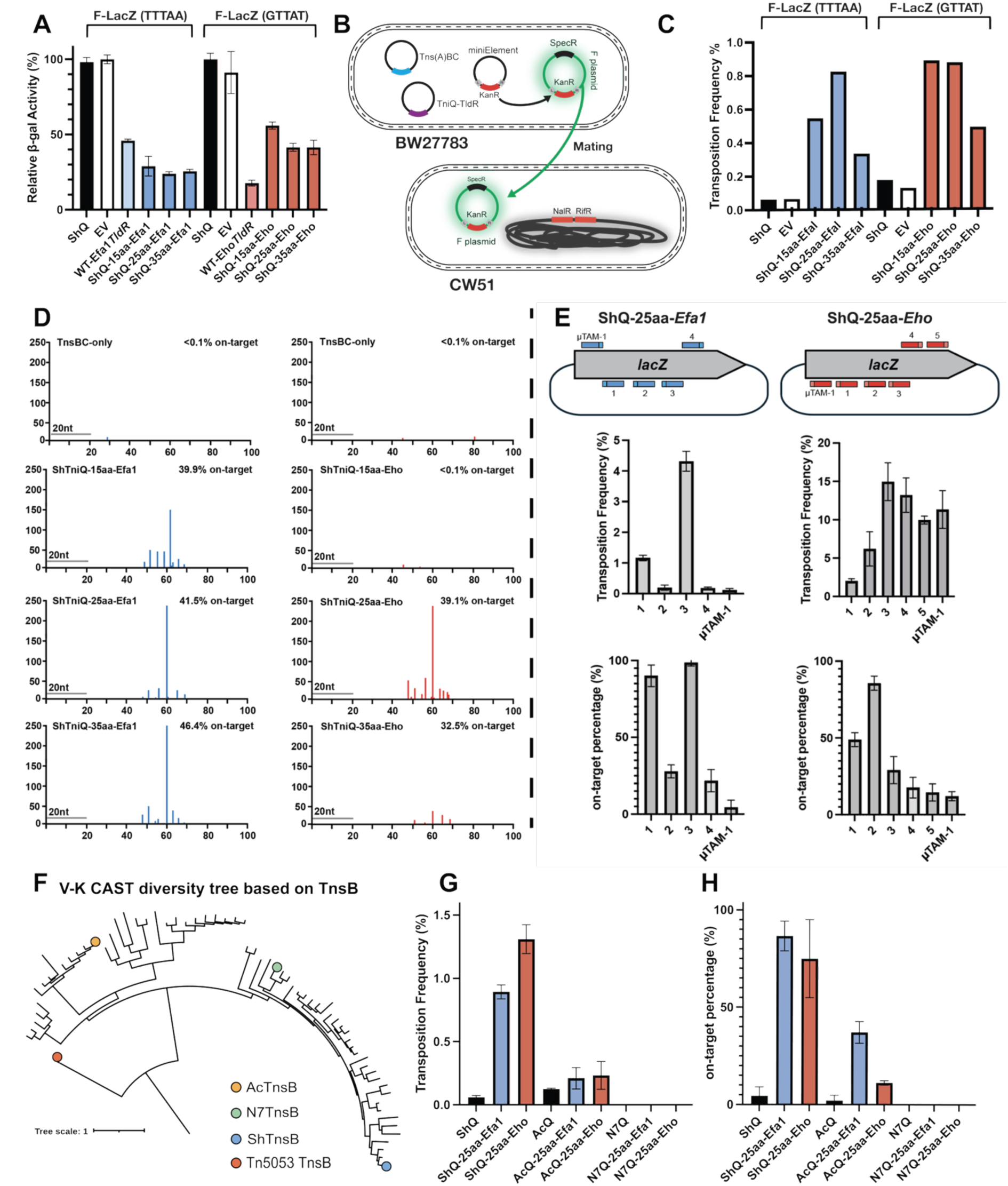
T**niQ-TldR fusions can effectively direct transposition by TnsB/C from type V-K CRISPR-associated transposon systems. A.** <u>β-gal</u> repression assay comparing wild-type TldRs and ShTniQ-TldR fusions ability to target P_lac_ and repress <u>β-gal</u> expression **B.** Mate out assay schematic. The donor strain contains all the necessary components for transposition. A donor element is targeted to an F-plasmid. F-plasmids are mated into recipient cells which are then plated on selective media. Transposition frequency is calculated by the ratio of kanamycin:spectinomycin-resistant colonies. **C.** Mate out assay using ShQ-TldRs to direct miniShDonor into F plasmid target. ShTniQ-EfaI and ShTniQ-Eho fusions boost transposition frequency across linker lengths compared to ShQ and EV controls. **D.** Illumina sequencing reads for ShTniQ-TldR fusions mapped to 100bp downstream of TAM (origin) on F-plasmid. Grey line and nucleotide label represent guideRNA length and binding site. On-site percentage is calculated by the insertions within the 47-68bp window downstream of the TAM relative to the total transposon reads in the F-plasmid. **E.** Transposition frequency and on-site percentage using different guides targeting *lacZ* on F plasmid. On-site frequency was determined by blue/white screening. Mutant TAMs (µTAMs) for Efa1 (WT-TTTAA; µ-ATTAA) and Eho (WT-GTTAT; µ-ATTAT) were tested alongside wild-type TAMs.

Given the finding that our fusion constructs could efficiently target DNA based on a genetic assay that monitors gene expression, we proceeded with testing their ability to target transposition using the previously described mating-out assay (**Figure 2B**).^5,33^ Compared to the TniQ-only and empty vector controls, we observed a higher transposition frequency in both TldRs and across all linker lengths (**Figure 2C**). To assess the ability of the system to produce on-site events, the F-plasmids that received a transposon were analyzed using Illumina DNA sequencing. Our sequencing indicated an enrichment of transposition within a ∼20bp window 47-68bp downstream of the TAM and target site (**Figure 2D**). This insertion window differs from the previously determined ShCAST window which is consistently 60-66bp from the PAM^2^. The wider insertion spacing in our TniQ-TldR system could result from the presumed lack of a direct protein-protein interaction between TniQ and the TldR that would stabilize the insertion window. While it is also possible that long linkers between ShTniQ and the TldR might contribute a wobble between the components that would widen the insertion window, longer linkers did not give a broader distribution than shorter linkers. Interestingly, in all cases, the largest insertion peak consistently resided 60-61bp from the TAM for both *Efa1*TldR and *Eho*TldR. Excluding TniQ-15aa-*Eho*, every fusion had a roughly 40% on-site insertion rate, which is similar to the 50-60% on-site insertion rate for wild-type V-K CAST.^2^ Additionally, insertions were only detected in one orientation with either fusion, indicating that our de novo system still functions unidirectionally.

To further characterize the ShTniQ-TldR fusions, we designed guide RNAs targeting multiple TAM-associated sites within *lacZ*. This also enabled us to pair the mating-out assay with blue/white screening to assess both transposition frequency and on-site insertion accuracy across conditions on selection plates with 5-Bromo-4-chloro-3-indolyl β-D-galactopyranoside (X-gal). For these and subsequent experiments, we used only the 25 amino acid linkers, as this length yielded consistently high on-site performance across our initial screens. We also evaluated the impact of TAM variants by designing guide sequences to target TAMs with an alternate nucleotide in the first position (e.g., TTTAA→ATTAA for *Efa1*TldR and GTTAT→ATTAT for *Eho*TldR). Transposition frequencies varied by as much as 20-fold across different guide RNAs for wild-type TAMs, and up to 40-fold with changes from the native TAMs (**Figure 2E**). Notably, we identified guides that yielded ∼99% on-site insertions for *Efa1*TldR and ∼86% for *Eho*TldR. Although the altered TAM supported high transposition frequency with *Eho*TldR, both TldRs failed to maintain targeting specificity under these conditions.

To test whether TniQ-TldR fusion functionality is specific to the ShCAST system or can extend to other closely related type V-K systems, we fused our TldRs to AcTniQ and N7TniQ derived from the *Anabaena cylindrica* (AcCAST) and *Nostoc sp*. PCC7101 (N7CAST) systems, respectively (**Figure 2F**). These two systems were selected because they are the best-characterized V-K CAST elements aside from ShCAST^2,14,59^. We found that the AcTniQ-TldR fusions could perform targeted transposition albeit at lower transposition frequencies and on-site transposition rates than the ShTniQ-TldR fusions (**Figure 2G-H**). Notably, despite N7CASTs closer relationship to ShCAST, we did not detect transposition for the N7TniQ-TldR fusions, suggesting that compatibility between TldRs and TniQ homologs is not solely determined by sequence similarity or evolutionary proximity.

### Alternate TniQ and effector fusion combinations fail to recapitulate gRNA-directed transposition

Encouraged by the results with TldR proteins fused to the V-K CAST-derived components, we next fused TldRs to TniQs from I-B1, I-D, and I-F3b CASTs. However, none of these combinations produced systems that were capable of gRNA-directed transposition (**Figure S2**). We also tested whether non-CAST Tn7-like elements could be converted into RNA-guided systems by fusing TldRs to the TniQ from the highly active Tn7-like transposon Tn6022, which lacks intrinsic sequence specificity (**Figure S2**).^29,30^ These fusions also failed to support targeted insertions. Additionally, we carried out a limited attempt to engineer a reduced I-F3 TniQ domain similar in size to the ShTniQ protein, but this also failed to produce a functional TldR-based guide RNA system (**Fig S2**).

To test whether the TnpB superfamily could more broadly serve as replacements for Cas12k, we fused ShTniQ to nuclease-deficient *D. radiodurans* TnpB (dTnpB) and *A. polyphaga* mimivirus Fanzor2 (dFz2) (**Figure S3A**). We chose a Fanzor2 representative due to their close structural and genetic similarity to prototypical TnpB, as opposed to the more diverged Fanzor1 branch.^25,26,31,32^ Unexpectedly, while we detected insertions for both TniQ-dTnpB and TniQ-dFz2, neither construct exhibited levels of on-site insertions above the TniQ-only and empty vector controls (**Figure S3B, C)**. Structurally, TldRs, TnpB,^23,34^ and Fanzor2^32^ proteins are similar, making it surprising that only TldRs could be successfully fused to ShTniQ. One possible explanation is that TldRs are adapted to bind DNA with high affinity as transcription factors, whereas TnpB and Fanzor2 proteins evolved primarily as nucleases. Stable DNA binding may be a unique feature of TldRs that is not shared among catalytically active members of the TnpB superfamily.

### Evaluating gRNA requirements for RNA-guided transposition

Since TldRs were originally characterized as RNA-guided transcriptional repressors, we suspected that discoveries related to their original biology might not directly translate to RNA-guided transposition, specifically in relation to the gRNA. Attributed to an unknown RNA-processing mechanism, *Efa1*TldR and *Eho*TldR were determined to support a maximum functional gRNA target length of 9nt and 16nt, respectively.^27^ These short guide sequence requirements are problematic in therapeutic contexts since they could lead to detrimental off-target insertions. To re-evaluate guide-length requirements in the context of RNA-guided transposition, we constructed gRNAs that were flanked by a 5′ hammerhead ribozyme and a 3′ hepatitis delta virus ribozyme, allowing for fixed-length guide sequences and autoprocessing of the overall gRNA scaffold (**Figure 3A**).^38,39^ Additionally, as is typical of gRNAs in the TnpB superfamily, the extended 5’ tail of TldRs was determined to be dispensable^27^, therefore we truncated our new TldR gRNA scaffold to ∼100nt, not including the guide sequence (**Figure 3A**). Using our mating-out assay, we observed an increase in transposition for the smaller gRNA with the 20nt fixed-length guide compared to the original larger gRNA with the 20-nt guide sequence, in both *Efa1*TldR and *Eho*TldR fusions (**Figure 3B**). Guide sequences with shorter lengths, including at the previously determined max-length (9 or 16nt), did not produce comparable levels of transposition frequency and on-site insertions as 20nt guide (**Figure 3B, C)**. Notably, the 9-nt guide for the *Efa1*TldR fusion yielded an on-site insertion rate of only 19%.

**Fig 3:**
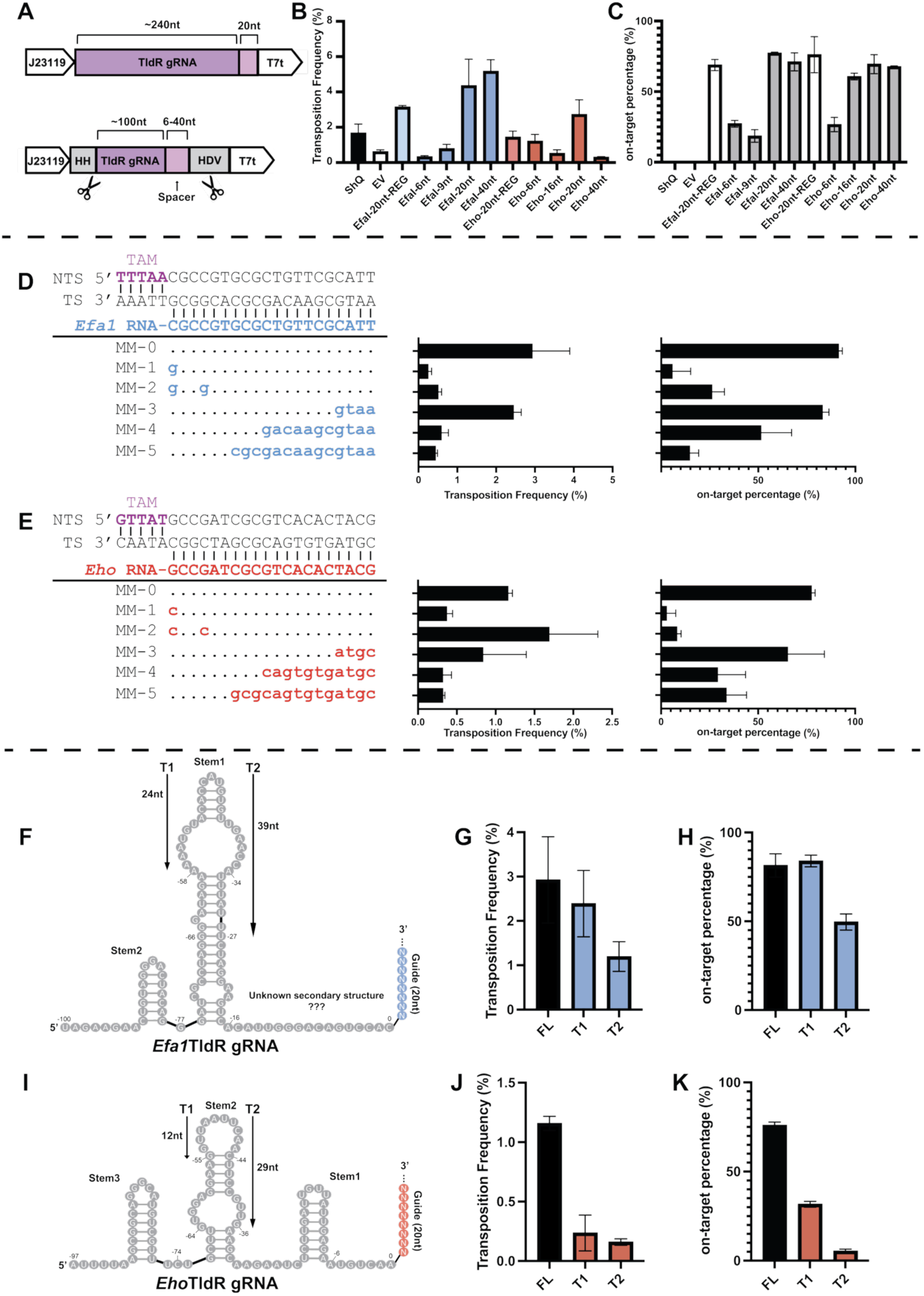
Guide RNA requirements for ShTniQ-TldR fusions. **A.** Diagram of original TldR-gRNA expression construct (left) versus preprocessed and minimized TldR-gRNA construct (right). Preprocessing is accomplished by expressing a hammerhead (HH) ribozyme at the 5’ end and a hepatitis delta virus (HDV) ribozyme at the 3’ end to produce a fixed guide length. **B-C.** Effect of gRNA preprocessing and guide length on transposition frequency (**B**) and on-site percentage (**C**) for TniQ-TldR fusions. The REG label refers to the original, unprocessed, unfixed 20nt gRNA expression construction. On-site frequency was determined by blue/white screening. **D-E.** Mismatch tolerance in ShQ-25aa-Efa1(**D**) and ShQ-25aa-Eho (**E**) measured by transposition frequency and on-site percentage. **F.** A 2D schematic of the *Efa1*TldR gRNA, showing the predicted secondary structure and sequence. The gRNA scaffold and guide region span nucleotides −100 to 0 and 1 to 21, respectively. 24nt and 39nt RNA truncations are labeled as T1 and T2, respectively. **G-H.** Transposition frequency (**G**) and on-site frequency (**H**) of *Efa1*TldR gRNA truncations compared to full-length (FL) gRNA. **I.** A 2D schematic of the *Eho*TldR gRNA, showing the predicted secondary structure and sequence. The gRNA scaffold and guide region span nucleotides −97 to 0 and 1 to 21, respectively. 12nt and 29nt RNA truncations are labeled as T1 and T2, respectively. **J-K.** Transposition frequency (**J**) and on-site frequency (**K**) of *Efa1*TldR gRNA truncations compared to full-length (FL) gRNA.

To further probe guide sequence requirements, we designed guides with varying degrees of RNA-DNA complementarity. Guide sequences with shorter complementary stretches again failed to support efficient transposition or specificity, even at 9nt and 16 nt lengths (**Figure 3D, E**). Moreover, minor mismatches in the TAM-proximal seed region significantly reduced on-site insertion percentages, similar to other TnpB superfamily proteins.^23,24,50–52^ Importantly, these results indicate that it is a 20nt length of sequence that is specific to the target that is important, not just the length of the guide RNA.

Together, these results suggest that a 20nt guide sequence functions, in the context of RNA-guided transposition, at a higher efficiency and specificity than the previously established shorter guides. While 9 and 16 nt guide sequences are sufficient for transcriptional repression, longer guide sequences appear to be more suitable for RNA-guided transposition. These findings also raise the possibility that 20 nt guide sequences could be broadly used to improve targeting precision in native TldRs, potentially minimizing off-site target effects if they are used for genome engineering applications.

We also examined the ability of fixed length autoprocessing gRNAs to function in the native ShCAST system. Contrasting our results with the ShQ-TldR fusions, we did not observe a significant boost in transposition frequency or on-site insertions (**Figure S4**). Because ShCas12k lacks intrinsic nuclease activity, it is thought to rely on host RNases to process its sgRNA. Thus, while auto-processing gRNAs did not enhance activity in the native host, they may still offer an advantage in heterologous systems where compatible bacterial RNases are absent.

Previous studies across the TnpB superfamily have shown that truncation of gRNA stem loops can increase activity and further minimize the system.^23,24,30,40^ To test this in our ShTniQ-TldR fusions, we designed several truncated gRNA variants for both *Efa1*TldR and *Eho*TldR and measured their transposition efficiencies. Due to the already compact size of their gRNAs, we chose to truncate only the largest stem loop, Stem2 (**Figure 3F, I**). We found that *Efa1*TldR’s gRNA tolerated truncation well, with the Trim1 variant maintaining comparable levels of on-site transposition (**Figure 3G, H**). However, the larger truncation (Trim2) reduced on-site activity to approximately 50%. In contrast, *Eho*TldR’s gRNA was less tolerant to truncation, with both Trim1 and Trim2 showing reduced transposition efficiency and lower on-site insertion rates (**Figure 3J, K**).

### ShTniQ-TldR fusions paired with nAniI-TnsB fusion can bypass the constraints of replicative transposition

Type V-K CAST systems transpose via replicative transposition, a process that should require recruiting a repair/restart DNA polymerase complex and that results in cointegrate insertions.^41^ In type I CASTs and in prototypical Tn7, the TnsA protein functions as a second essential transposase component that cleaves the non-transferred strand just inside the donor DNA thereby enabling cut-and-paste transposition.^3,34,35^ Previous studies with ShCAST have shown that fusing ShTnsB to I-nAniI, a homing nicking endonuclease, enables cut-and-paste transposition (**Figure 4A**).^14^ To test for cointegrate formation and resolution in the TldR-based system, we introduced a tetracycline resistance marker into the donor plasmid backbone. This allowed us to monitor insertion of the backbone during transposition using the mating-out assay. By comparing the number of kanamycin-only colonies to those with both kanamycin and tetracycline resistance, we could determine the percentage of transposition events that were retained as cointegrates in the assay. We found that pairing TniQ-TldR fusions with nAniI-TnsB fusions virtually eliminated transposition products indicating cointegrate formation while retaining high on-site insertion rates (**Figure 4B-D**). Specifically, our mating-out assay showed that, compared to TnsB alone, the nAniI-TnsB fusion reduced cointegrate formation from ∼40% to <4% for both the *Efa1*TldR and *Eho*TldR ShTniQ fusions.

**Fig 4:**
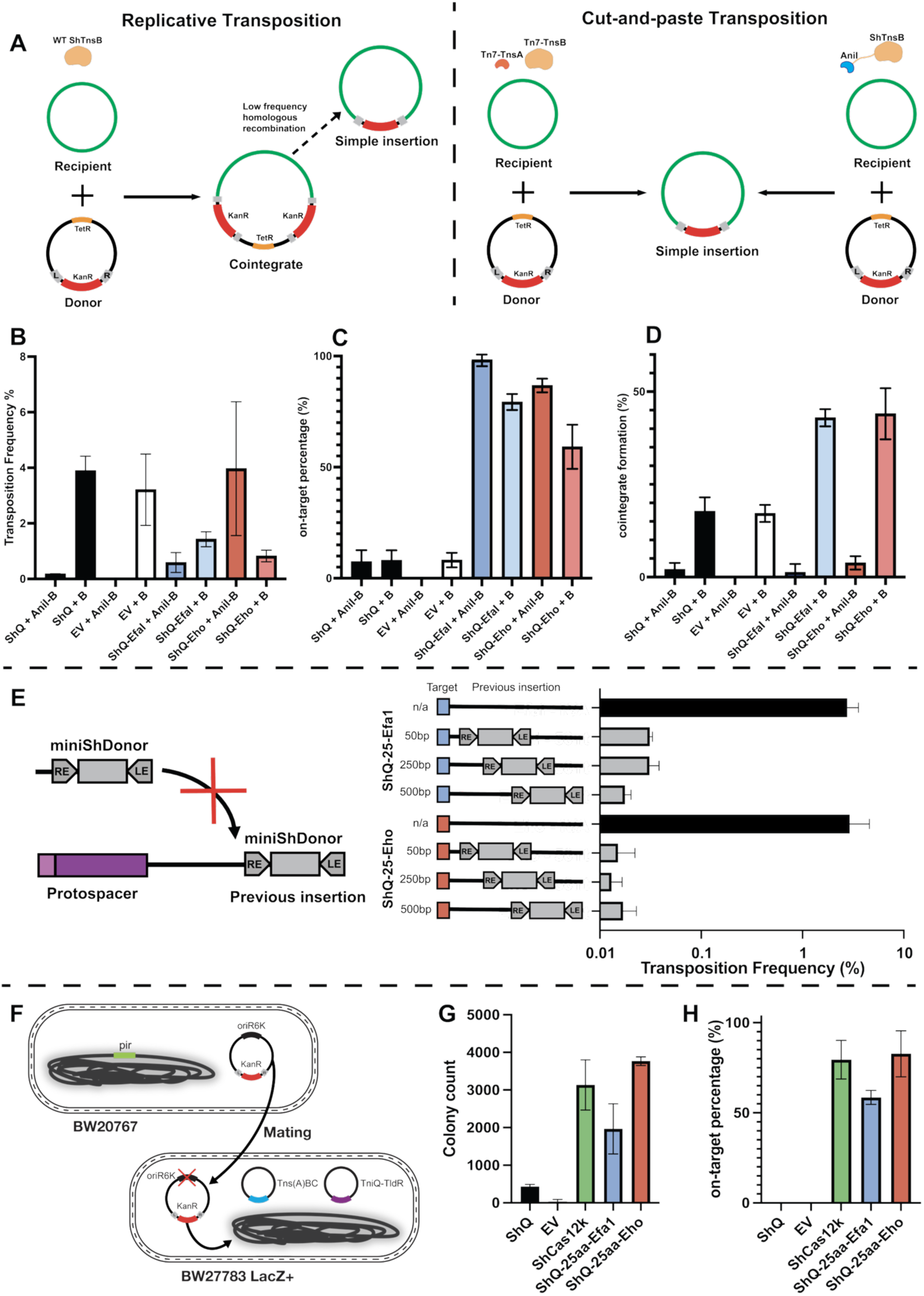
Demonstration of Cut-and-Paste Activity, Target Immunity, and Chromosomal Insertions in TniQ–TldR Transposition. **A.** Schematics comparing transposition mechanisms resulting in simple or cointegrate insertions in wild-type ShTnsB, engineered nAniI-ShTnsB, and TnsA/B from Tn7. **B-D.** Transposition frequency (**B)**, on-site percentage (**C**), and cointegrate formation (**D**) of ShTniQ-TldR fusions paired with nAniI-TnsB fusions. On-site frequency was determined by blue/white screening. **E.** Effect of previously inserted ShDonor element at varying distances from target on ShTniQ-TldR transposition frequency. **F.** Mating-in assay schematic. The donor strain contains the donor element on a plasmid with a R6K replicon. The recipient strain contains all the components for transposition and does not support replication of the donor element plasmid. The donor and recipient strains are mated followed by being plated on selective media. Integration in the genome results in successful growth on selective media. **G-H.** Colony count (**G**) and on-site frequency (**H**) for mating-assay.

### The de novo TldR gRNA transposition system retains target immunity

A hallmark of prototypical Tn7 is its ability to prevent multiple insertions from occurring at the same targeted site.^36,37,44^ Whether this feature would be recapitulated after redirecting a Tn7-like system with an entirely different targeting module was unknown. To evaluate this, we designed four different F plasmid targets for each ShTniQ-TldR. Three targets were constructed that had mini elements positioned 50, 250, and 500 bp from the programmed integration site in the target which were compared to one without a copy of the element already residing proximal to the integration site. We observed a strong inhibitory effect on transposition at all tested distances, with nearly a 100-fold decrease compared to the control (**Figure 4E**), indicating that this crucial regulatory feature of Tn7-like elements is recapitulated in the reprogrammed systems. This study represents the first demonstration of redirecting a Tn7-like system with non-CRISPR components, raising the question of whether the molecular safeguards that prevent reinsertion would remain functional. That the TldR integrations systems possess target immunity is an important finding for potential therapeutic applications. It was critical that ShTniQ-TldR fusions retain this feature, as uncontrolled multiple insertions at a single locus could be harmful.

### de novo TldR transposition system can efficiently transpose into the bacterial chromosome

To evaluate whether the ShTniQ-TldR fusions can mediate RNA-guided integration into the bacterial chromosome and to compare their efficiency to WT ShCas12k, we used the previously described mating-in assay^5^. This assay differs from our standard mating-out setup, which measures integration into an F plasmid target, by testing the ability of the system to mobilize and integrate DNA cargo into the recipient cell’s genome (**Figure 4F**). In this assay, donor strains carrying the transposon mini-element were mated with recipient *E. coli* strains harboring the appropriate TldR fusion and transposition machinery. Following conjugation and antibiotic selection, transconjugants were screened for insertion events at chromosomal *lacZ*. We also included X-gal in the selection medium, to visually assess on-site insertion frequency via blue/white screening.

We observed a boost in transposition for both ShTniQ-TldR fusions compared to the ShTniQ-only and empty vector controls and were able to confirm the ability of our fusion construct to target transposition into the chromosome (**Figure 4G-H**). The ShTniQ-*Efa1* and ShTniQ-*Eho* fusions achieved roughly 58% and 83% on-site insertion frequency, respectively (**Figure 4H**). This efficiency rivals that of ShCas12k in which roughly 79% of insertions were on target. These results establish that ShTniQ-TldR fusions are capable of mediating efficient and specific RNA-guided integration into the bacterial genome at rates comparable to WT V-K CAST systems.

### Genetically attenuating off-site targeting in the ShCAST system

One of the initial limitations identified in early work with the V-K CAST systems was a substantial amount of off-site targeting events that resulted from effector-independent transposition. Off-site nuclease activity will have serious negative consequences in canonical CRISPR-Cas systems imposing DNA damage even in the absence of an invading bacteriophage, plasmid or other genetic elements. However, a low level of transposition that does not require a specific guide RNA would presumably not have the same negative consequence and could be advantageous for the host and the transposon, allowing an additional avenue for new integration sites even without a gRNA.

The structure of ShCAST transpososome^10,11^ reveals that TniQ bridges the interaction between the TnsC filament and the Cas12K-sgRNA section. We hypothesized that weakening or otherwise altering the binding interface between TniQ and TnsC might make the system more dependent on the Cas12K effector complex and thereby enhance on-site transposition. TniQ interacts with the two terminal protomers (11^th^ and 12^th^) of the TnsC filament (**Figure 5A**), occluding a surface area of almost 900 Å^2^ with protomer 11^th^ and 300 Å^2^ with protomer 12^th^. The interaction with protomer 11^th^ is mediated by the N-terminal region of TniQ (residues Met1-Leu13) as well as the loop containing Asn33 and His34, while the interaction with the terminal TnsC protomer is mediated by residue Glu147-Asp148 and Arg155 (**Figure 5A-C**).

**Fig 5:**
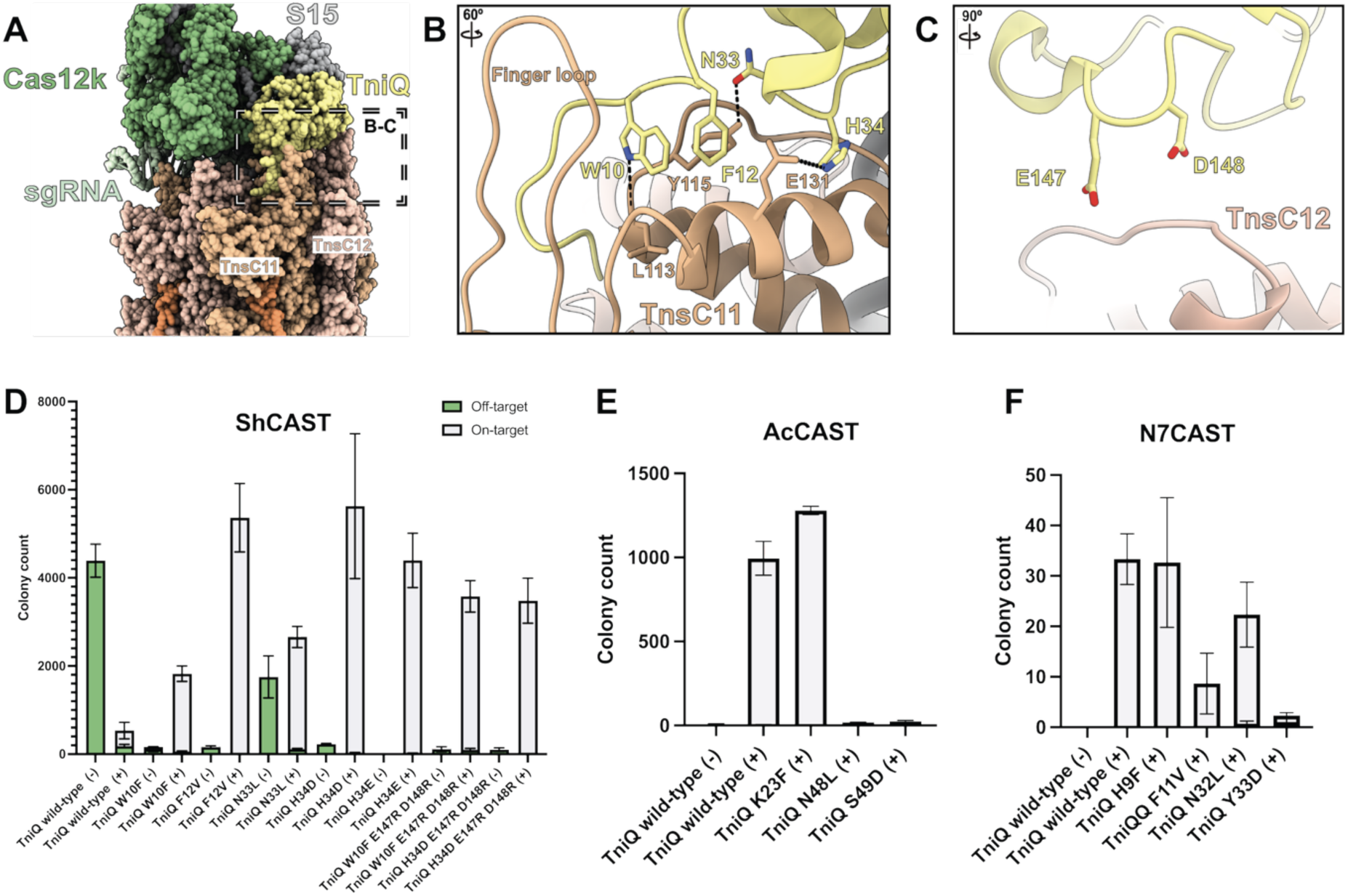
Modifications in the TniQ-TnsC interface improve the performance of ShCAST system. **A.** Partial view of the structure of ShCAST transpososome where the TniQ(yellow)-TnsC(light and dark orange; indicating different subunits) interface is highlighted with a black and white dotted square. Hydrogen bond interactions are represented by black dashed lines. **B.** Zoomed in view of the TniQ-TnsC interface, specifically the interaction of TniQ with subunit 11 of TnsC. **C.** Zoomed in view of the TnsC-TniQ interface, specifically the interaction of TniQ with subunit 12 of TnsC. **D.** Transposition frequency of mutated TniQ variants from ShCAST system. For each TniQ mutant, two bars are shown: one for untargeted transposition using BCQ only (–), and one for targeted transposition using BCQ + sgRNA + Cas12k (+). The identity of each TniQ mutation is indicated on the x-axis. Transposition frequency was measured by blue/white screening using sgRNA_MP2 that targets lacZ. **E.** Transposition frequency of mutated TniQ variants from AcCAST system. **F.** Transposition frequency of mutated TniQ variants from N7CAST system.

We first explored the defects associated with altering the TniQ interaction with the 11^th^ TnsC protomer. Previous studies have shown that deletion of the first twelve amino acids of TniQ (Met1-Phe12) completely abrogates transposition.^11^ Within this region, mutating Trp10 (TniQ-W10A) had a more severe transposition defect than deleting the first eight residues of TniQ, suggesting that Trp10 and Phe12 anchor the interaction between the N-terminus of TniQ and the 11^th^ protomer of the TnsC filament. Therefore, we generated variants of TniQ with conservative mutations at these positions. The TniQ-W10F and TniQ-F12V variants improved ShCAST performance by reducing the amount of off-site transposition and increasing the overall transposition rate in presence of Cas12k and sgRNA. Point mutations on the second loop mediating the interaction with this protomer of the filament (N33L and H34D/E) had comparable effects (**Figure 5D, S5**). Interestingly, pairing mutations in these two loops results in lower to null transposition frequencies (**Figure S5**), suggesting that the Cas12K effector can only overcome subtle defects on this TniQ-TnsC interface. Mutating the residues that interact with protomer 12^th^ of the TnsC filament affected only mildly the rate of on-site transposition and overall transposition frequency, and combinations with the W10F and H34D mutations yielded similar results to those mutations alone (**Figure 5D, S5**). A consistent trend across variants was that the reduction of effector-independent transposition often coincided with an increase in effector-dependent transposition (**Figure 5D**). Collectively, these results show that Cas12k and sgRNA can bypass minor defects on the TniQ-TnsC interface providing an avenue to enhance on-site transposition.

We selected some of the best performing TniQ mutant variants from ShCAST and tested equivalent mutations in two other V-K CAST systems, AcCAST and N7CAST, using the same spacer as for ShCAST (**Figure S6**). Our first observation was that, at least in our experimental set-up, the rate of untargeted transposition for AcCAST and N7CAST was very low, while the on-site frequency was higher compared to ShCAST, despite using the same spacer. Notably, AcCAST exhibited an approximate 10-fold higher frequency of transposition compared to ShCAST and N7CAST. Differences may have been expected given that AcCAST is phylogenetically more diverged from the other two systems (**Figure 2F**) The mutations in TniQ that improved the performance of ShCAST had either no effect on transposition frequency or in other cases negatively affected AcCAST and N7CAST transposition. One possible explanation is that the mutations selected in ShCAST may not map to functionally equivalent positions in the AcCAST and N7CAST TniQ proteins. While we chose corresponding residues based on sequence alignment, it is possible that mutations with similar functional effects occur at different sites in AcCAST and N7CAST, given the divergence observed in the TniQ alignment.

These data suggest the TniQ and TnsC interface provides a critical balance for establishing productive transposition that is still sensitive to signals imparted by the effector. Small changes along this interface will likely drive adaptation for elements to display more or less strict on- and off-site targeting. It is possible that the various type V-K CAST are under different selective pressures in the native environment where they naturally exist. For example, the environment for AcCAST and N7CAST may be more selective for precise targeting while some random transposition may be more beneficial in the natural environment for the ShCAST element. Alternatively, differences in the intrinsic activity or regulation of the associated Cas12k proteins, or variation in DNA-binding kinetics, may also contribute to the observed transposition profiles across systems.

The same mutations that reduced off-site targeting in the ShCAST system did not have a noticeable effect with the ShTniQ-TldR systems (**Fig S9**). This is consistent with our Alphafold3 model showing that there is not a direct interaction between the TldR and TniQ that could compensate for a reduced TniQ-TnsC interaction (**Figure 1D**).

It was also noted previously that the addition of the nAniI nuclease domain to TnsB reduced the amount of ShCAST off-site targeting that was found with transposition,^14^ something that we also observed in our work (**Figure S7**). We suspect that the addition of a new domain to the transposon component subtly changes the ratio of activities of components and changes in this balance effect on- and off-site targeting. This has been proposed previously with a focus on the amount of TnsC expression compared to other components.^47^ In our own work, we have also found that fusing GFP domains to TnsC is permissible and also alters the on-verses off-site targeting found in these assays (**Figure S8**). All these observations suggest that optimizing the ratio of transposition components will be an important consideration when adapting the type V-K-based system to any new editable host.

## Discussion

We have demonstrated that TniQ-TldR fusions can be used to engineer de novo, RNA-guided transposition systems that rivals natural V-K CAST in specificity and orientation control, while substantially reducing coding burden and reliance on a host factor. By varying guide length, processing, and transposase configuration, we reveal both the engineering potential and the biology of TldR effectors repurposed for DNA integration.

Our data shows alternate requirements for TldR-guided transposition as compared to their roles as stand-alone repressors from previous work. Whereas native TldRs were reported to repress transcription with only 9-16 nt of guide complementarity,^27^ we find that a full 20 nt guide maximizes both transposition frequency and on-target precision (**Figure 3B,C)**. Mutations even in the distal region of the 20 nt guide significantly reduced insertion efficiency suggesting that extending complementarity beyond 16 nt can markedly suppress off-target events. Incorporation of 5′ HH and 3′ HDV ribozymes to create a fixed-length gRNA further boosts activity, potentially alleviating requirement of host RNA processing enzymes.

Both *Efa1*TldR and *Eho*TldR require 5 nucleotide TAMs which could limit the pool of suitable targets in a host, however previous studies in TnpB and Fanzor2 variants have shown rational mutagenesis approaches can relax TAM constraints without compromising activity and precision.^48,49^ Future studies will need to be conducted to introduce TAM promiscuity and will likely be applicable to TldRs broadly as an RNA-guided DNA binding protein.

This work represents the first example of an engineered TnpB-family protein as a module for programmable DNA binding. Although TnpBs and Fanzors share the same bilobed core and ancestry,^23,24^ only TldRs fused to TniQ yielded robust, programmable transposition. Though AlphaFold3 predictions of TldR versus TnpB/Fz2 do not reveal obvious structural differences to explain this functional divergence (**Figure S1**). The failure of catalytically dead TnpB or Fz2 fusions could suggest that TldRs possess uniquely favorable DNA-binding kinetics or RNP stability. Currently, no studies have shown whether the guide target sequence activity is conserved across TnpB-family proteins. Furthermore, guide-walking experiments could be performed with dTnpB and dFz2 to potentially find a more suitable gRNA. Further biochemical experiments will need to be conducted to compare the DNA-binding kinetics between TnpBs, TldRs, and Fanzors.

To investigate whether TniQ-TldR fusion-mediated transposition is a unique feature of the ShCAST system or can be generalized across diverse Tn7-like elements, we constructed and tested fusions between our TldR proteins and TniQ homologs derived from I-B1, I-D, and I-F3b CAST systems, as well as the non-CRISPR Tn6022 transposon. As mentioned previously, we did not observe targeted transposition in any of these alternate TniQ-TldR fusions. These findings align with at least one example showing that a dCas9 fusion to TniQ from a I-F3a CAST was not able to execute targeted transposition.^3^ This suggests that the V-K family of CAST systems may possess unique structural or mechanistic features that make them particularly amenable to our fusion strategy. One potential explanation lies in the size difference: V-K CAST TniQs are unusually small (∼170–190 amino acids) compared to those in most Tn7-like systems, which typically range from 300 to 500 amino acids. Supporting this idea, we observed successful targeted transposition when fusing TldRs to AcTniQ (184aa) from the V-K CAST family. However, fusions involving N7TniQ (167 amino acids), a V-K CAST family member more closely related to ShTniQ, did not yield detectable transposition activity. Clearly additional factors beyond size and evolutionary lineage influence fusion compatibility such as subtle structural differences, mechanistic differences, or protein expression levels. Further structural and biochemical characterization will be important to elucidate these determinants and may enable the rational design of fusion constructs applicable across broader Tn7-like transposon systems.

In parallel, we explored an alternative strategy to improve the specificity of ShCAST by introducing targeted mutations at the TniQ-TnsC interface. One of the trends that was identified was that multiple changes in one of the predicted contact subregions between TniQ and TnsC often compromised transposition activity. The greatest level of improvement with increasing on-siteing and reducing off-site targeting was achieved by single changes across all three predicted points of contact between TniQ and the terminal and subterminal protomers of TnsC. Also of note, changes meant to be synonymous in the AcCAST and N7CAST systems did not show the same improvements in on-siteing but instead often compromised transposition activity. AcCAST and N7CAST also showed better on-siteing in their native state when tested in our *E. coli* assay system, something that could indicate that this interaction surface was sufficiently weak that further perturbations would not increase a dependance on the effector complex for driving stability. Taken together this work suggests that the TniQ-TnsC interface is a critical contact point for setting the balance between on- and off-site targeting. The previous observation that the relative level of TnsC expression to other components, and the effect of additional protein domains (i.e. nAniI and GFP) alters on- and off-site transposition could also support the importance of this balance. Further optimization of natural systems will likely benefit from comparing many different examples and optimizing the stability of the transpososome in the final target host to be genetically edited. Alternatively, the de novo strategy we pursued in this work could offer more engineering potential that is not constrained by the natural environment where these systems evolved.

We expect that future work with more TldRs and TnpBs will allow further optimization of these important guide RNA directed transposition systems. We refer to these engineered systems as Repressor Operated Unidirectional Transposable Elements (ROUTEs), highlighting the use of repressor-derived effectors and its preservation of directional integration behavior. Together, these results demonstrate that TldRs can be repurposed as programmable DNA-targeting modules for RNA-guided transposition. Although TldRs are evolutionarily derived from the TnpB superfamily, they are not naturally associated with canonical mobile genetic elements or possess CRISPR arrays. As such, the ROUTE system represents a de novo class of RNA-guided transposons, expanding the design space of Tn7-like elements beyond CRISPR-associated systems. More broadly, these findings suggest that TldRs could serve as compact and programmable targeting modules for a wide range of RNA-guided applications, including base-editors, epigenome editors, transcriptional activators, and fluorescent tags.

## Acknowledgments

This work was funded with a grant from the National Institutes of Health to JEP, GM152260. RDS was funded by the National Science Foundation Graduate Research Fellowship Program, DGE – 2139899.

## Declaration of Interests

Cornell University has submitted patent applications for results in this manuscript where some of the authors are listed as inventors.

## Methods

### Growth conditions

*Escherichia coli* strains (Table S1) were grown in lysogeny broth (LB) or on LB agar supplemented with the following concentrations of antibiotics when appropriate: 100 μg/ml carbenicillin, 30 μg/ml chloramphenicol, 20 μg/ml tetracycline, 50 μg/ml kanamycin, 50 μg/ml spectinomycin, 20 μg/ml nalidixic acid, 100 μg/ml rifampicin, 75 μg/ml X-gal.

### β-galactosidase Repression Assay

DNA-binding activity was tested via a modified version of the previously described Miller Assay. Vectors encoding the fusions constructs were transformed into BW27783^57^ cells containing a modified F plasmid with *lacZ* and the corresponding target sequence on P*_lac_*. Overnight cultures were grown in LB medium supplemented with appropriate antibiotics at 37°C. Cultures were diluted 1:25 into fresh LB and grown to mid-log phase (OD₆₀₀ ∼0.3–0.5). Cultures were washed and resuspended in LB supplemented with 1 mM IPTG and or 0.2% (w/v) L-arabinose. Cells were incubated for an additional 2 hours at 37°C with shaking. Cultures were harvested and transferred to a clean tube. OD₆₀₀ was measured using a spectrophotometer and recorded for normalization. A 1mL aliquot of each culture was permeabilized by addition of 50 µL 0.1% SDS and 200 µL chloroform. To initiate the reaction, 20 µL of culture was incubated in 600 <u>µL of</u> 60 mM Na₂HPO₄, 40 mM NaH₂PO₄, 5 mM β-mercaptoethanol, and 1 mg/mL ONPG for 90 minutes at <u>30°C</u>. Reactions were terminated by the addition of 700 µL of 1 M Na₂CO₃. Samples were centrifuged briefly to remove debris, and absorbance at 420 nm (A₄₂₀) was measured. Miller Units were calculated using the standard formula.

### Mating-out transposition assays

Transposition frequency was assessed in a large pool of independent transformants, based on a previous assay.^5^ Briefly, vectors encoding the core transposition machinery (TnsABC or TnsBC) and our fusion construct (TniQ-TldR) were co-transformed into BW27783^57^ cells containing a modified F plasmid with a corresponding target sequence and separate donor plasmid containing the mini-element (Kanamycin resistance gene flanked by left and right transposon ends). Plates were grown overnight, and hundreds of transformants were washed off the plate in M9 minimal media, pelleted, washed twice with M9 minimal media, and finally resuspended to O.D. 0.6 in M9 minimal media supplemented with required antibiotics, 0.2% w/v maltose, 0.2% w/v arabinose, and 0.1 mM IPTG. After 20 h of incubation with shaking at 37°C, 0.5 ml of the donor cells was spun down, washed twice with LB, and resuspended into 0.5 ml LB supplemented with 0.2% w/v glucose for recovery with shaking at 37°C for 1.5 hours. To monitor transposition from the donor plasmid into the F plasmid target, donor cells were then mixed with mid-log recipient cells (CW51^33^) in LB supplemented with 0.2% w/v glucose at a ratio of 1:5 donor:recipient and incubated with gentle agitation for 90 minutes at 37°C to allow mating. After incubation, cultures were vortexed, placed on ice, then serially diluted in LB 0.2% w/v glucose and plated on LB supplemented with required antibiotics for selecting CW51 recipient cells for transconjugants nalidixic acid, rifampicin, spectinomycin, with or without kanamycin to sample the entire transconjugant population or select for transposition respectively. Plates were incubated at 37°C for 24 h before colonies were counted. To test for on/off-site target frequency, plates were supplemented with X-gal.

In mating-out experiments monitoring cointegrate formation, a modified assay was performed in which the donor plasmid contained a tetracycline resistance gene, in addition to the kanamycin gene within the mini-element. After the final mating-out assay incubation, cultures were plated on LB supplemented with nalidixic acid, rifampicin, and spectinomycin. Including the aforementioned antibiotics, cultures were also plated on kanamycin or kanamycin+tetracycline to select for overall transposition and cointegrate transposition, respectively.

### Mapping insertions

Illumina sequencing was used to map the total insertions from F plasmids from transconjugants. Transconjugants were pooled, diluted 1/100 in fresh LB containing kanamycin and grown to mid-log. F plasmid DNA was isolated using the ZR BAC DNA Miniprep Kit. Insertions were mapped with BBtools (BBMap – Bushnell B. – sourceforge.net/projects/bbmap/).

### Mating-in transposition assays

Transposition into the *E. coli* chromsome was monitored via a mate-in transposition assay, as previously described.^6^ To make the donor strain, a donor plasmid carrying a mini-element element and a conditional origin of replication was transformed into BW20767.^58^ The recipient strain PO619^6^ (*Escherichia coli* BW27783 *lacZYA*^+^) was created by transforming vectors carrying core transposition machinery. Single colonies from each strain were grown overnight in LB media at 37°C. Overnight recipient cultures were diluted 1:50 in LB induction media (0.1mM IPTG, 0.2% (w/v) arabinose, 30 μg/mL chloramphenicol, 50 μg/mL carbenicillin). Overnight donor cultures were diluted 1:25 in LB supplemented with 50 μg/mL Kanamycin. Both sets of cultures were grown to mid-log. Cultures of donor and recipient strains were spun down, washed with LB twice, and resuspended to O.D.600 = 10. The donor cells were mixed with recipients in a ratio of 1:5, cells were spun down and resuspended in LB supplemented with 0.1 mM IPTG and 0.2% (w/v) arabinose. Conjugal mating was performed at 37°C for 2 h. After mating, cells were spun down and plated on LB plates supplemented with the carbenicillin, chloramphenicol, and kanamycin. To test for on/off-site target frequency, plates were supplemented with X-gal.

### TnsB similarity tree construction

Annotated genomic sequences and feature tables of Cyanobacteria were downloaded from National Center for Biotechnology Information (NCBI) FTP site. There were 6580 annotated genomes when sequences were downloaded for our analysis (June 2025). The HMM profiles associated with TnsB (PF06527) and Cas12K (NF038191) downloaded from the European Bioinformatics Institute (EMBL-EBI) Pfam database, were used for detecting homologs with hmmsearch (HMMER3). Only TnsB hits that had a close Cas12K hit -30 kb or less apart in the same genomic file-were selected for further analysis. Entries marked as ‘partial’ or with a length lower than 450 amino acids were removed. Proteins with more than 90% identity were clustered together using cd-hit. The curated protein list, with the addition of TnsB from Tn5053, was aligned using MUSCLE^53^ and a similarity tree was constructed using FastTree.^54^ The tree was visualized in iTOL,^55^ using TnsB from Tn5053 as the outgroup.

## Figures

**Fig S1:**
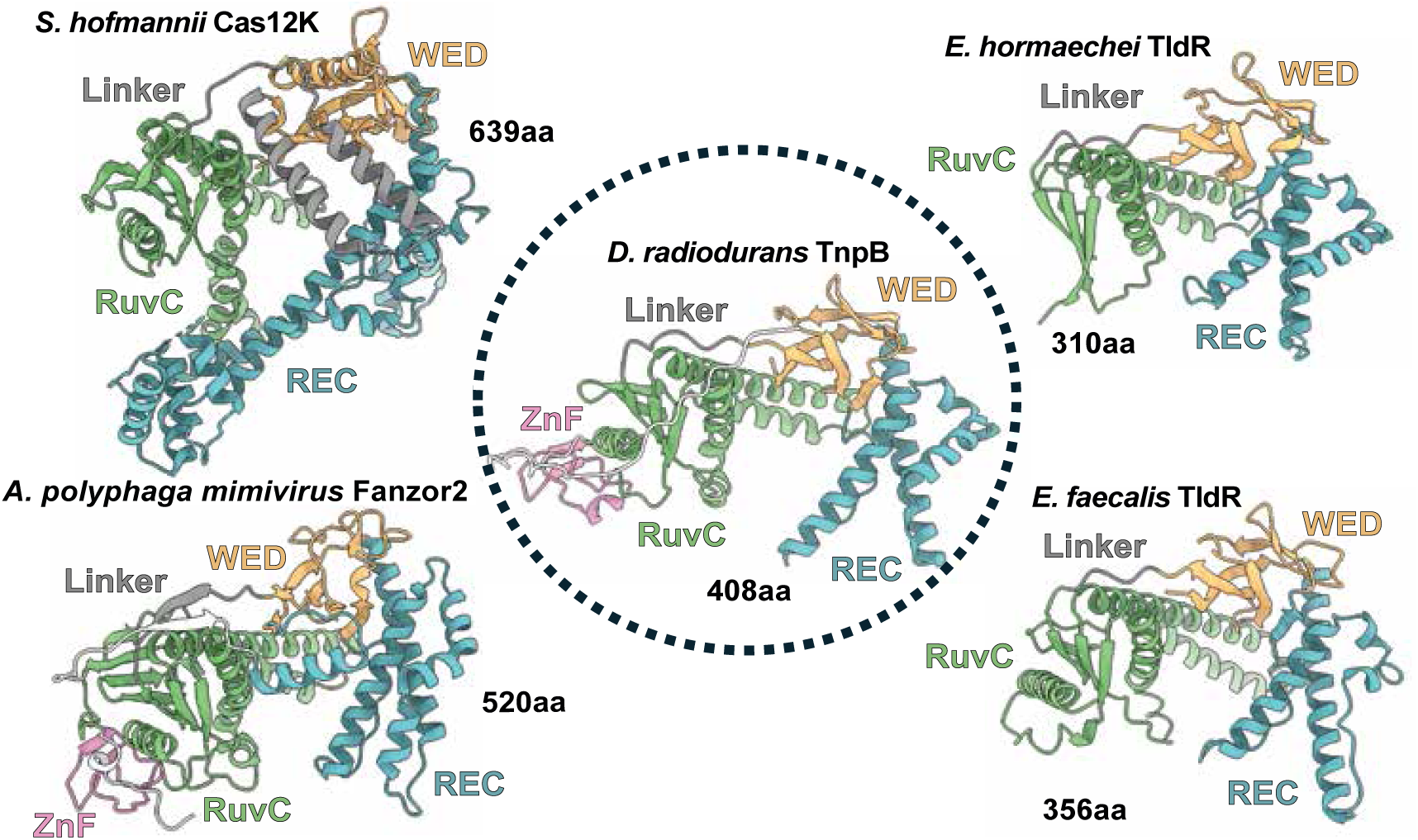
Structural comparison of TnpB-family representatives. Architectural comparison between TnpB and descendant proteins. *D. radiodurans* (middle; 8H1J)^24^, *S. hofmannii* Cas12k (top left; 8EA3)^10^, and *A. polyphaga* mimivirus Fanzor2 (bottom left; 9B0L)^31^ structures were determined by cryoEM in previous studies. *E. faecalis* (top right) and *E. hormaechei* (bottom right) structures were determined by AlphaFold3^28^. Cas12k and TldRs lack the ZnF domain rendering them catalytically dead.

**Fig S2:**
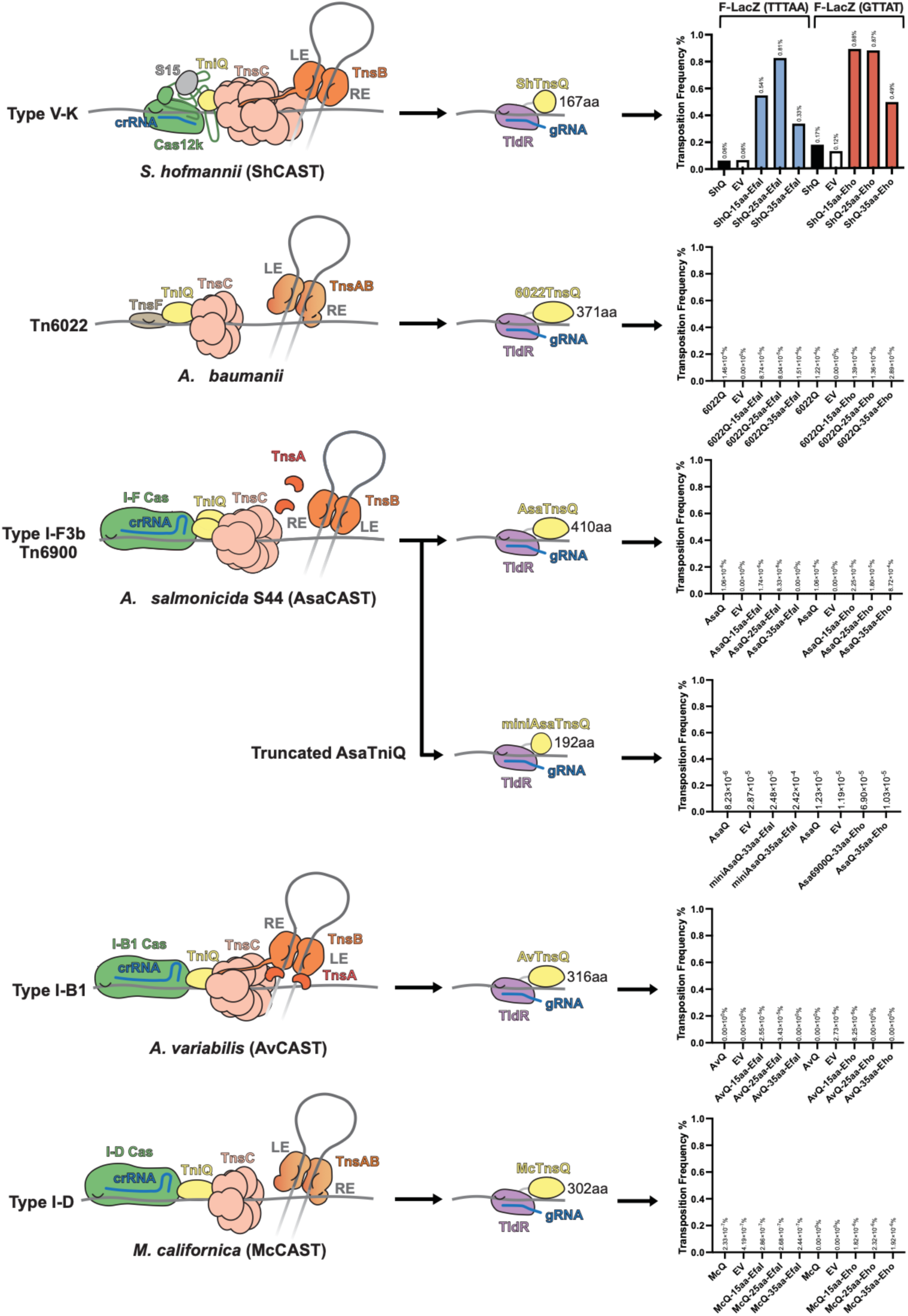
TniQs from various Tn7-like systems fail to stimulate transposition when fused to TldRs. Mate out assay using various TniQs from CRISPR-associated and non-CRISPR-associated Tn7-like elements to direct transposition into F plasmid target. Only ShTniQ from V-K CAST can be fused to TldRs to stimulate transposition.

**Fig S3:**
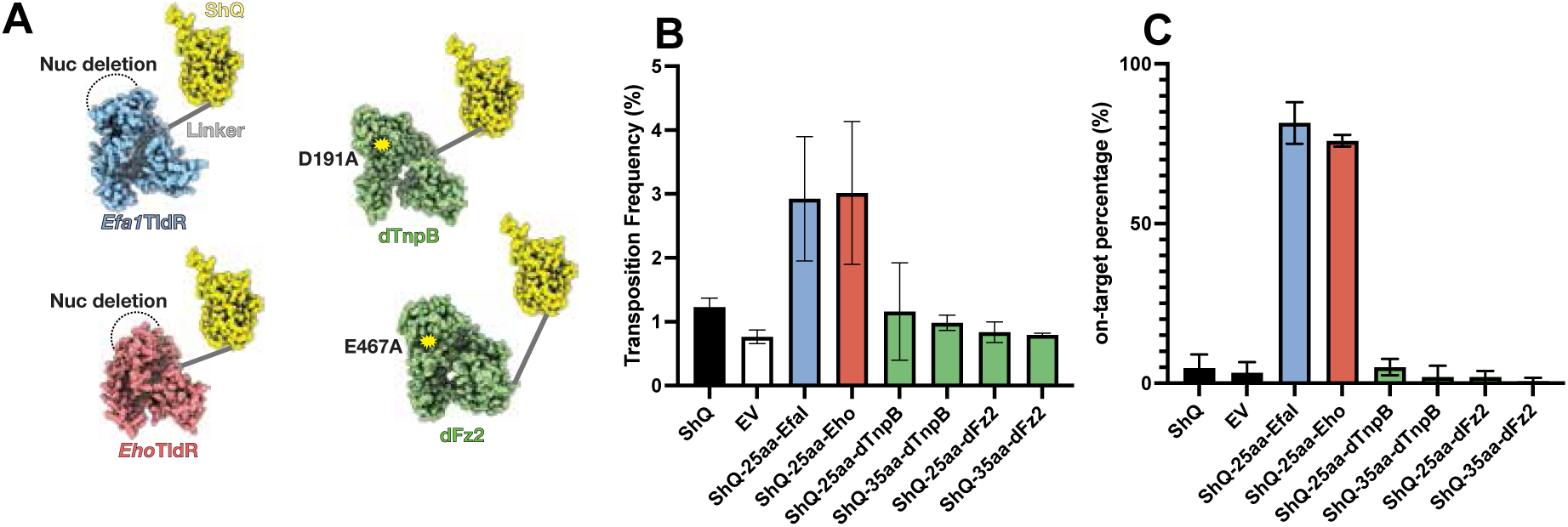
ShTniQ fusions to dTnpB or dFz2 do not stimulate targeted transposition. **A.** Diagram showing representations of Efa1, Eho, dTnpB, and dFanzor2 fused to ShTniQ. Dra2TnpB and ApmFanzor2’s nuclease activity was inactivated using D191A and E467A mutants, respectively. **B-C.** Transposition Frequency (**B)** and on-site percentage (**C**) compared between Efa1 (targeting LacZ2, shown in Figure 2E; left) and Eho (targeting LacZ3, shown in Figure 2E; right). Despite close structural similarity and evolutionary lineage, ShTniQ fusions to dTnpB and dFz2 do not stimulate targeted transposition.

**Fig S4:**
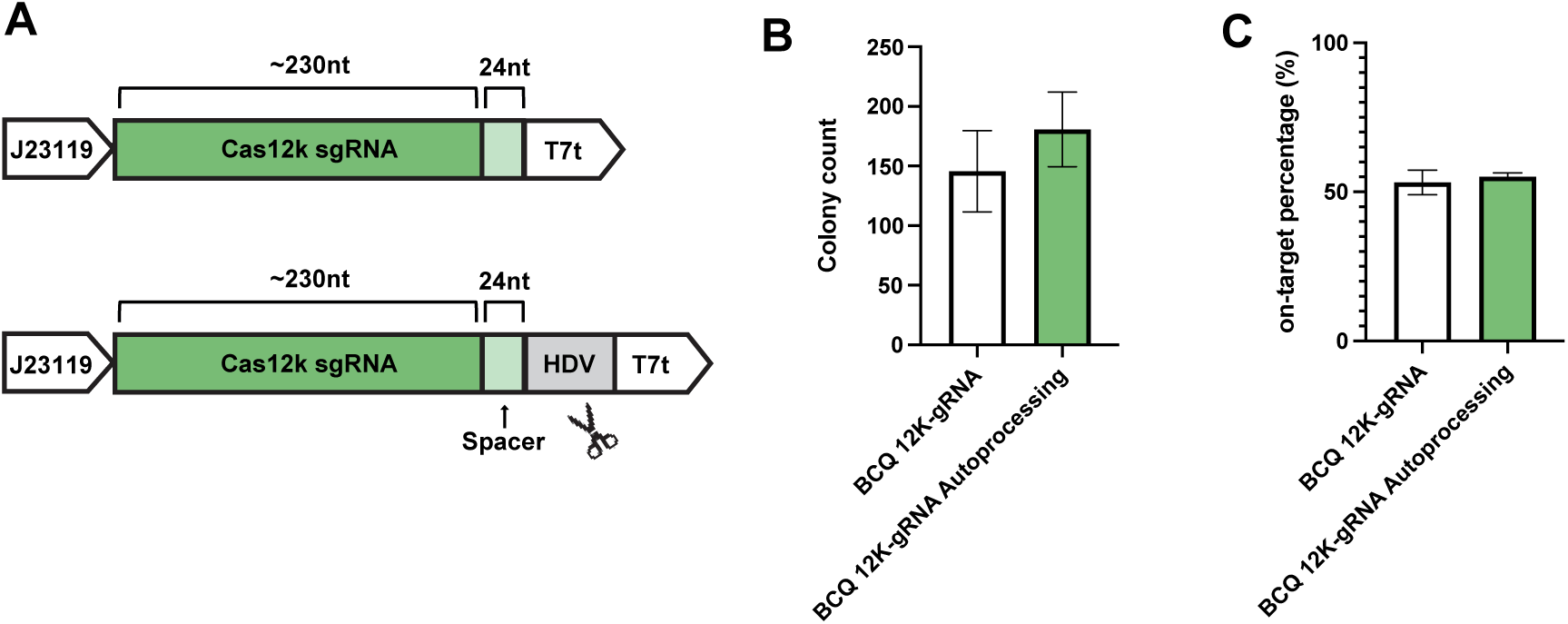
Effect of fixed-length guideRNA on wild-type ShCAST system. **A.** Diagram of original Cas12k-gRNA expression construct (top) versus fixed-length construct (bottom). Preprocessing is accomplished by expressing a hepatitis delta virus (HDV) ribozyme at the 3’ end to produce a fixed guide length. **B-C.** Effect of gRNA preprocessing on transposition frequency (**B**) and on-site percentage (**C**) for wild-type ShCAST system.

**Fig S5:**
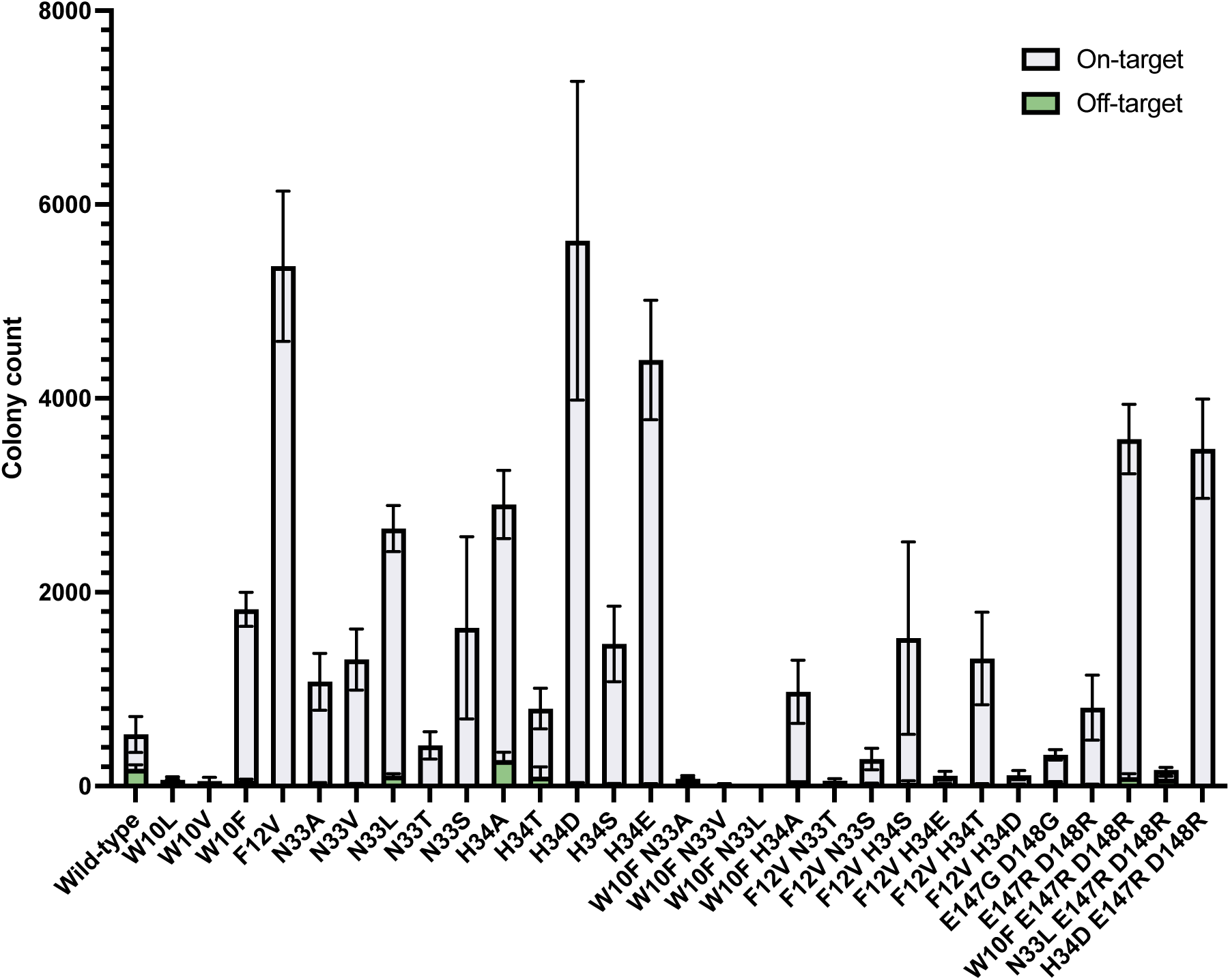
Transposition frequency of ShCAST using mutated TniQ variants. Each bar corresponds to a different TniQ mutant. All measurements correspond to targeted transposition using BCQ + sgRNA + Cas12k. The identity of each TniQ mutation is indicated on the x-axis. Transposition frequency was measured by blue/white screening using sgRNA-MP2 that targets *lacZ*.

**Fig S6:**
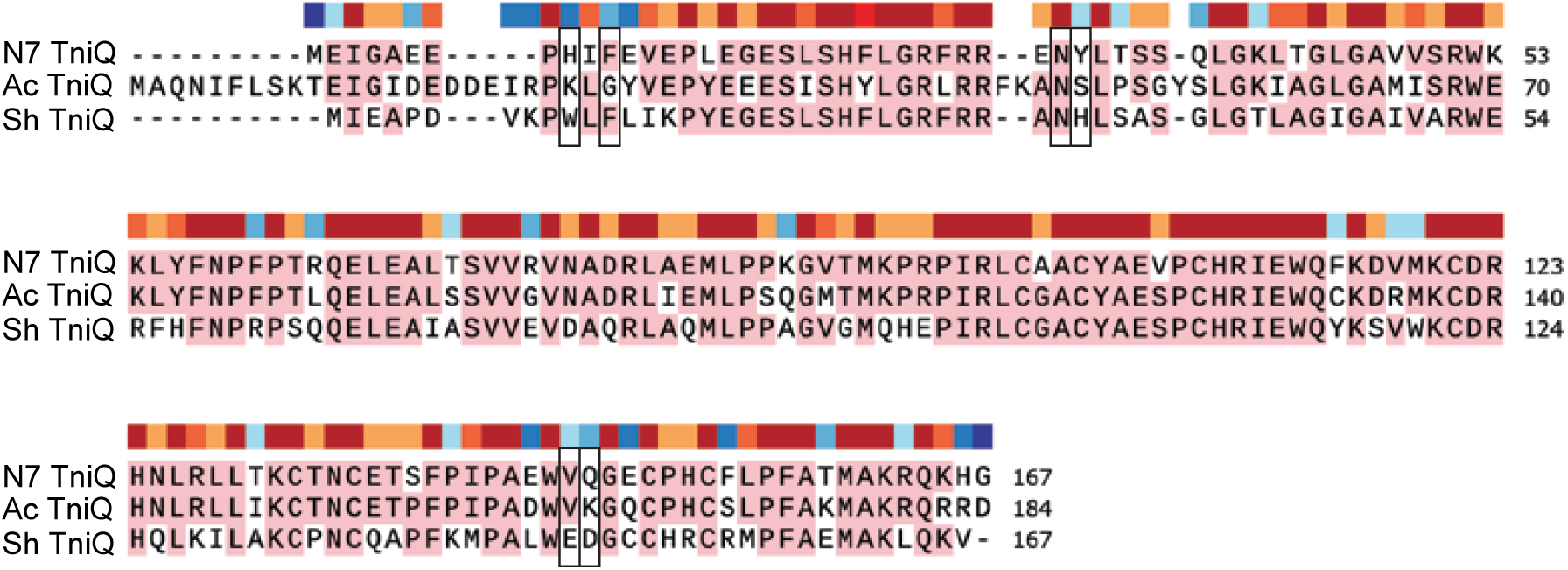
Protein sequence alignment of TniQ from type V-K CAST. Alignment of TniQ proteins from Ac, N7 and ShCAST systems generated using MUSCLE (accessed via SnapGene). The colored bars above the alignment indicate the degree of residue conservation, with a gradient from blue (highly variable) to red (highly conserved). Identical residues across all three sequences are highlighted in pink. Positions subjected to mutagenesis in ShCAST and their predicted equivalents in the other two systems are indicated by a black square in the alignment.

**Fig S7:**
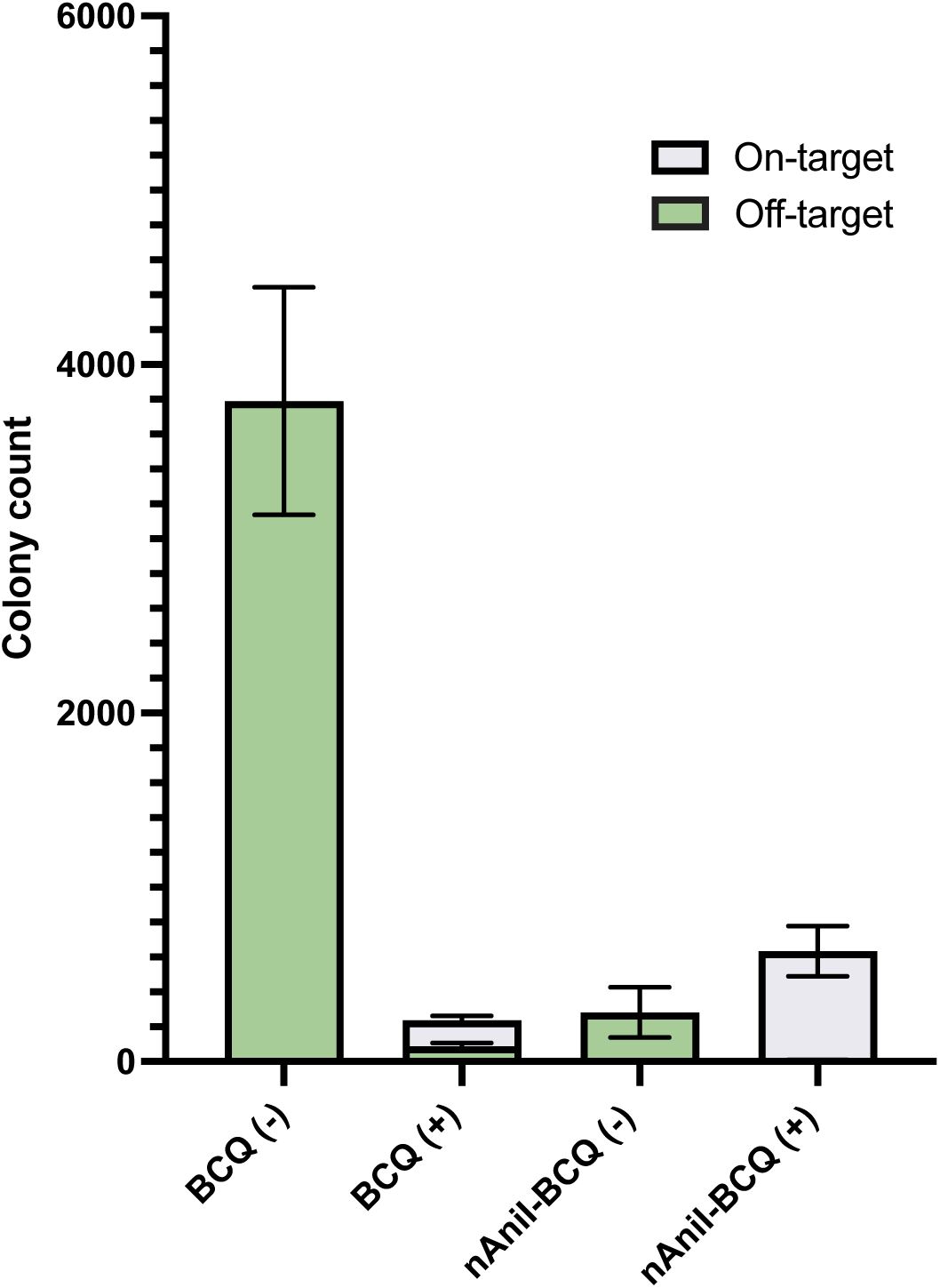
Effect of fusing nAniI to TnsB in ShCAST. Transposition frequency of ShCAST with nAniI-TnsB fusion. For wild-type and nAniI-fused TnsB, two bars are shown: one for untargeted transposition using BCQ only (-), and one for targeted transposition using BCQ + sgRNA + Cas12k (+). Transposition frequency was measured by blue/white screening using sgRNA_MP2 that targets *lacZ*.

**Fig S8:**
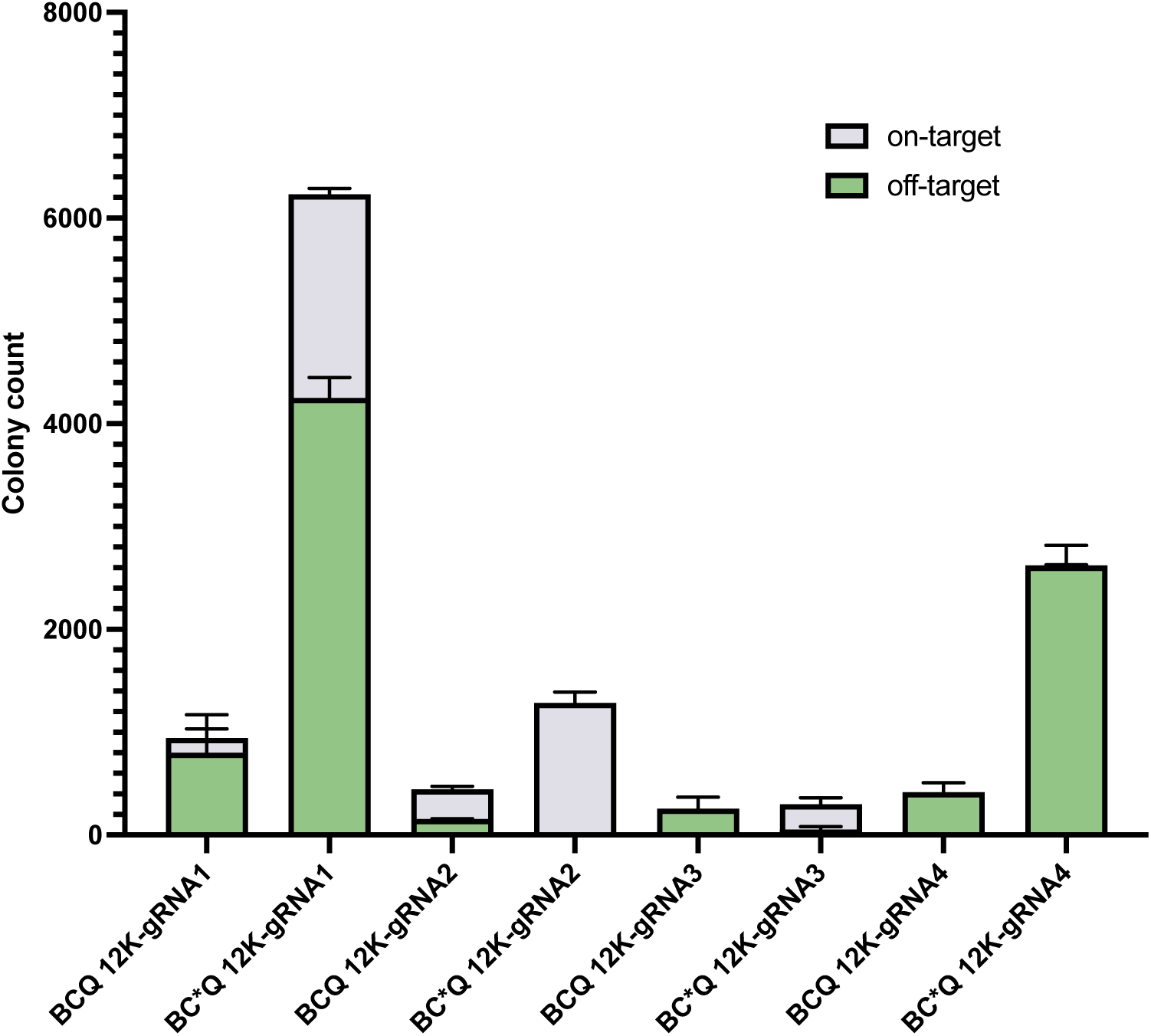
Effect of TnsC-GFP fusions on targeted transposition in ShCAST. Transposition frequency of TnsC-GFP fusions using four different gRNAs targeting *lacZ*. All conditions reflect targeted transposition using BCQ + sgRNA + Cas12k. The graph compares transposition frequencies achieved with wild-type TnsC (BCQ) versus a TnsC-GFP fusion protein (denoted as BC*Q on the x-axis) across four different guide RNAs (gRNA1–gRNA4). Transposition frequency was measured by blue/white screening and is presented as colony counts, with on-site events shown in light gray and off-target events in cyan. Data represents the mean ± SD of biological replicates.

**Fig S9:**
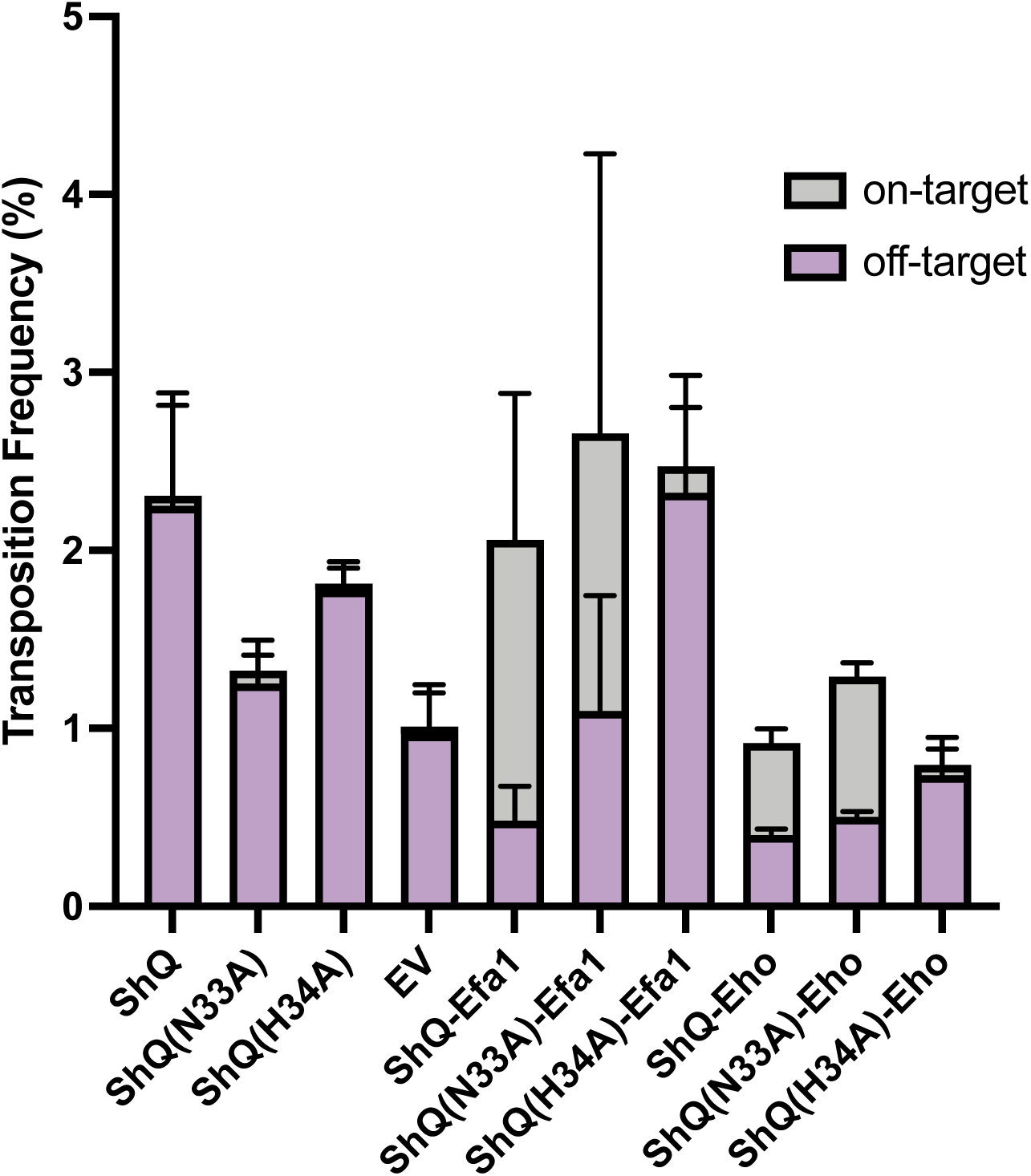
Assaying ShTniQ mutants in the TniQ-TldR fusion construct. Transposition and on-site frequency of ShTniQ mutants in the TniQ-TldR construct. On-site frequency was measured by blue/white screening, with on-site events shown in light gray and off-target events in purple. Data represents the mean ± SD of biological replicates.

## References

1. Peters, J.E., Makarova, K.S., Shmakov, S., and Koonin, E.V. (2017). Recruitment of CRISPR-cas systems by TN7-like transposons. Proceedings of the National Academy of Sciences 114.

2. Strecker, J., Ladha, A., Gardner, Z., Schmid-Burgk, J.L., Makarova, K.S., Koonin, E.V., and Zhang, F. (2019). RNA-guided DNA insertion with CRISPR-associated transposases. Science 365, 48–53.

3. Klompe, S.E., Vo, P.L., Halpin-Healy, T.S., and Sternberg, S.H. (2019). Transposon-encoded CRISPR–CAS systems direct RNA-guided DNA integration. Nature 571, 219– 225.

4. Faure, G., Shmakov, S.A., Yan, W.X., Cheng, D.R., Scott, D.A., Peters, J.E., Makarova, K.S., and Koonin, E.V. (2019). CRISPR–Cas in mobile genetic elements: Counter-defence and beyond. Nature Reviews Microbiology 17, 513–525.

5. Petassi, M.T., Hsieh, S.-C., and Peters, J.E. (2020). Guide RNA categorization enables target site choice in TN7-CRISPR-Cas Transposons. Cell 183.

6. Hsieh, S.-C., and Peters, J.E. (2023). Discovery and characterization of novel type I-D CRISPR-guided transposons identified among diverse TN7-like elements in cyanobacteria. Nucleic Acids Research 51, 765–782.

7. Saito, M., Ladha, A., Strecker, J., Faure, G., Neumann, E., Altae-Tran, H., Macrae, R.K., and Zhang, F. (2021). Dual modes of CRISPR-associated transposon homing. Cell 184.

8. Hsieh, S.-C., and Peters, J.E. (2024). Natural and Engineered Guide RNA–directed transposition with CRISPR-associated TN7-like transposons. Annual Review of Biochemistry 93, 139–161.

9. Querques, I., Schmitz, M., Oberli, S., Chanez, C., and Jinek, M. (2021). Target site selection and remodelling by type V CRISPR-Transposon Systems. Nature 599, 497–502.

10. Park, J.-U., Tsai, A.W.-L., Rizo, A.N., Truong, V.H., Wellner, T.X., Schargel, R.D., and Kellogg, E.H. (2022). Structures of the HOLO CRISPR RNA-guided Transposon Integration Complex. Nature 613, 775–782.

11. Schmitz, M., Querques, I., Oberli, S., Chanez, C., and Jinek, M. (2022). Structural basis for the Assembly of the Type V CRISPR-associated transposon complex. Cell 185.

12. Park, J.-U., Tsai, A.W.-L., Mehrotra, E., Petassi, M.T., Hsieh, S.-C., Ke, A., Peters, J.E., and Kellogg, E.H. (2021). Structural basis for target site selection in RNA-guided DNA transposition systems. Science 373, 768–774. 10.1126/science.abi8976.

13. Park, J.-U., Tsai, A.W.-L., Chen, T.H., Peters, J.E., and Kellogg, E.H. (2022). Mechanistic details of CRISPR-associated transposon recruitment and integration revealed by cryo-EM. Proceedings of the National Academy of Sciences 119. 10.1073/pnas.2202590119.

14. Tou, C.J., Orr, B., and Kleinstiver, B.P. (2023). Precise cut-and-paste DNA insertion using engineered type V-K CRISPR-associated transposases. Nature Biotechnology 41, 968– 979. 10.1038/s41587-022-01574-x.

15. Hsieh, S.-C., and Peters, J.E. (2024). Natural and Engineered Guide RNA–Directed Transposition with CRISPR-Associated Tn7-Like Transposons. Annual Review of Biochemistry 93, 139–161. 10.1146/annurev-biochem-030122-041908.

16. Zeng, T., Yin, J., Liu, Z., Li, Z., Zhang, Y., Lv, Y., Lu, M.-L., Luo, M., Chen, M., and Xiao, Y. (2023). Mechanistic insights into transposon cleavage and integration by TnsB of ShCAST system. Cell Reports 42, 112698. 10.1016/j.celrep.2023.112698.

17. Altae-Tran, H., Shmakov, S.A., Makarova, K.S., Wolf, Y.I., Kannan, S., Zhang, F., and Koonin, E.V. (2023). Diversity, evolution, and classification of the RNA-guided nucleases TnpB and Cas12. Proceedings of the National Academy of Sciences 120. 10.1073/pnas.2308224120.

18. Siguier, P., Gourbeyre, E., and Chandler, M. (2014). Bacterial insertion sequences: their genomic impact and diversity. FEMS Microbiology Reviews 38, 865–891. 10.1111/1574-6976.12067.

19. Shmakov, S., Smargon, A., Scott, D., Cox, D., Pyzocha, N., Yan, W., Abudayyeh, O.O., Gootenberg, J.S., Makarova, K.S., Wolf, Y.I., et al. (2017). Diversity and evolution of class 2 CRISPR–Cas systems. Nature Reviews Microbiology 15, 169–182. 10.1038/nrmicro.2016.184.

20. Bao, W., and Jurka, J. (2013). Homologues of bacterial TnpB_IS605 are widespread in diverse eukaryotic transposable elements. Mobile DNA 4, 12. 10.1186/1759-8753-4-12.

21. Karvelis, T., Druteika, G., Bigelyte, G., Budre, K., Zedaveinyte, R., Silanskas, A., Kazlauskas, D., Venclovas, Č., and Siksnys, V. (2021). Transposon-associated TnpB is a programmable RNA-guided DNA endonuclease. Nature 599, 692–696. 10.1038/s41586-021-04058-1.

22. Altae-Tran, H., Kannan, S., Demircioglu, F.E., Oshiro, R., Nety, S.P., McKay, L.J., Dlakić, M., Inskeep, W.P., Makarova, K.S., Macrae, R.K., et al. (2021). The widespread IS200/IS605 transposon family encodes diverse programmable RNA-guided endonucleases. Science 374, 57–65. 10.1126/science.abj6856.

23. Nakagawa, R., Hirano, H., Omura, S.N., Nety, S., Kannan, S., Altae-Tran, H., Yao, X., Sakaguchi, Y., Ohira, T., Wu, W.Y., et al. (2023). Cryo-EM structure of the transposon-associated TnpB enzyme. Nature 616, 390–397. 10.1038/s41586-023-05933-9.

24. Sasnauskas, G., Tamulaitiene, G., Druteika, G., Carabias, A., Silanskas, A., Kazlauskas, D., Venclovas, Č., Montoya, G., Karvelis, T., and Siksnys, V. (2023). TnpB structure reveals minimal functional core of Cas12 nuclease family. Nature 616, 384–389. 10.1038/s41586-023-05826-x.

25. Saito, M., Xu, P., Faure, G., Maguire, S., Kannan, S., Altae-Tran, H., Vo, S., Desimone, A., Macrae, R.K., and Zhang, F. (2023). Fanzor is a eukaryotic programmable RNA-guided endonuclease. Nature 620, 660–668. 10.1038/s41586-023-06356-2.

26. Jiang, K., Lim, J., Sgrizzi, S., Trinh, M., Kayabolen, A., Yutin, N., Bao, W., Kato, K., Koonin, E.V., Gootenberg, J.S., et al. (2023). Programmable RNA-guided DNA endonucleases are widespread in eukaryotes and their viruses. Science Advances 9. 10.1126/sciadv.adk0171.

27. Wiegand, T., Hoffmann, F.T., Walker, M.W.G., Tang, S., Richard, E., Le, H.C., Meers, C., and Sternberg, S.H. (2024). TnpB homologues exapted from transposons are RNA-guided transcription factors. Nature 631, 439–448. 10.1038/s41586-024-07598-4.

28. Abramson, J., Adler, J., Dunger, J., Evans, R., Green, T., Pritzel, A., Ronneberger, O., Willmore, L., Ballard, A.J., Bambrick, J., et al. (2024). Accurate structure prediction of biomolecular interactions with AlphaFold 3. Nature 630, 493–500. 10.1038/s41586-024-07487-w.

29. Faure, G., Saito, M., Benler, S., Peng, I., Wolf, Y.I., Strecker, J., Altae-Tran, H., Neumann, E., Li, D., Makarova, K.S., et al. (2023). Modularity and diversity of target selectors in Tn7 transposons. Molecular Cell 83, 2122–2136.e10. 10.1016/j.molcel.2023.05.013.

30. Correa, A., Shehreen, S., Machado, L.C., Thesier, J., Cunic, L.M., Petassi, M.T., Chu, J., Kapili, B.J., Jia, Y., England, K.A., et al. (2024). Novel mechanisms of diversity generation in Acinetobacter baumannii resistance islands driven by Tn7-like elements. Nucleic Acids Research 52, 3180–3198. 10.1093/nar/gkae129.

31. Schargel, R.D., Qayyum, M.Z., Tanwar, A.S., Kalathur, R.C., and Kellogg, E.H. (2024). Structure of Fanzor2 reveals insights into the evolution of the TnpB superfamily. Nature Structural & Molecular Biology. 10.1038/s41594-024-01394-4.

32. Yoon, P.H., Skopintsev, P., Shi, H., Chen, L., Adler, B.A., Al-Shimary, M., Craig, R.J., Loi, K.J., DeTurk, E.C., Li, Z., et al. (2023). Eukaryotic RNA-guided endonucleases evolved from a unique clade of bacterial enzymes. Nucleic Acids Research 51, 12414–12427. 10.1093/nar/gkad1053.

33. Waddell, C.S., and Craig, N.L. (1988). Tn7 transposition: two transposition pathways directed by five Tn7-encoded genes. Genes & Development 2, 137–149. 10.1101/gad.2.2.137.

34. Peters, J.E., and Craig, N.L. (2001). Tn7: smarter than we thought. Nature Reviews Molecular Cell Biology 2, 806–814. 10.1038/35099006.

35. Peters, J.E. (2014). TN7. Microbiology Spectrum 2. 10.1128/microbiolspec.mdna3-0010-2014.

36. Skelding, Z. (2003). Alternative interactions between the Tn7 transposase and the Tn7 target DNA binding protein regulate target immunity and transposition. The EMBO Journal 22, 5904–5917. 10.1093/emboj/cdg551.

37. Stellwagen, A.E. (1997). Avoiding self: two Tn7-encoded proteins mediate target immunity in Tn7 transposition. The EMBO Journal 16, 6823–6834. 10.1093/emboj/16.22.6823.

38. Berkhout, B., Gao, Z., and Herrera-Carrillo, E. (2020). Design and Evaluation of Guide RNA Transcripts with a 3′-Terminal HDV Ribozyme to Enhance CRISPR-Based Gene Inactivation. Methods in Molecular Biology, 205–224. 10.1007/978-1-0716-0716-9_12.

39. Gao, Z., Herrera-Carrillo, E., and Berkhout, B. (2018). Improvement of the CRISPR-Cpf1 system with ribozyme-processed crRNA. RNA Biology 15, 1458–1467. 10.1080/15476286.2018.1551703.

40. Thornton, B.W., Weissman, R.F., Tran, R.V., Duong, B.T., Rodriguez, J.E., Terrace, C.I., Groover, E.D., Park, J.-U., Tartaglia, J., Doudna, J.A., et al. (2025). Discovery of widespread activating mutations in a compact RNA-guided endonuclease. bioRxiv (Cold Spring Harbor Laboratory). 10.1101/2025.02.11.637750.

41. Rice, P.A., Craig, N.L., and Dyda, F. (2020). Comment on “RNA-guided DNA insertion with CRISPR-associated transposases.” Science 368. 10.1126/science.abb2022.

42. Jang, S., Sandler, S.J., and Harshey, R.M. (2012). Mu Insertions Are Repaired by the Double-Strand Break Repair Pathway of Escherichia coli. PLoS Genetics 8, e1002642. 10.1371/journal.pgen.1002642.

43. Choi, W., Jang, S., and Harshey, R.M. (2014). Mu transpososome and RecBCD nuclease collaborate in the repair of simple Mu insertions. Proceedings of the National Academy of Sciences 111, 14112–14117. 10.1073/pnas.1407562111.

44. DeBoy, R.T., and Craig, N.L. (1996). Tn7 transposition as a probe of cis interactions between widely separated (190 kilobases apart) DNA sites in the Escherichia coli chromosome. Journal of Bacteriology 178, 6184–6191. 10.1128/jb.178.21.6184-6191.1996.

45. Hsieh, S.-C., Fülöp, M., Schargel, R., Petassi, M.T., Barabas, O., and Peters, J.E. (2025). Telomeric transposons are pervasive in linear bacterial genomes. Science. 10.1126/science.adp1973.

46. Wu, W.Y., Adiego-Pérez, B., and Van Der Oost, J. (2024). Biology and applications of CRISPR–Cas12 and transposon-associated homologs. Nature Biotechnology. 10.1038/s41587-024-02485-9.

47. George, J.T., Acree, C., Park, J.-U., Kong, M., Wiegand, T., Pignot, Y.L., Kellogg, E.H., Greene, E.C., and Sternberg, S.H. (2023). Mechanism of target site selection by type V-K CRISPR-associated transposases. Science 382. 10.1126/science.adj8543.

48. Marquart, K.F., Mathis, N., Mollaysa, A., Müller, S., Kissling, L., Rothgangl, T., Schmidheini, L., Kulcsár, P.I., Allam, A., Kaufmann, M.M., et al. (2024). Effective genome editing with an enhanced ISDra2 TnpB system and deep learning-predicted ωRNAs. Nature Methods. 10.1038/s41592-024-02418-z.

49. Wei, Y., Gao, P., Pan, D., Li, G., Chen, Y., Li, S., Jiang, H., Yue, Y., Wu, Z., Liu, Z., et al. (2025). Engineering eukaryotic transposon-encoded Fanzor2 system for genome editing in mammals. Nature Chemical Biology. 10.1038/s41589-025-01902-7.

50. Swarts, D.C., Van Der Oost, J., and Jinek, M. (2017). Structural basis for guide RNA processing and Seed-Dependent DNA targeting by CRISPR-CAS12A. Molecular Cell 66, 221–233.e4. 10.1016/j.molcel.2017.03.016.

51. Jain, I., Minakhin, L., Mekler, V., Sitnik, V., Rubanova, N., Severinov, K., and Semenova, E. (2018). Defining the seed sequence of the Cas12b CRISPR-Cas effector complex. RNA Biology 16, 413–422. 10.1080/15476286.2018.1495492.

52. Pausch, P., Soczek, K.M., Herbst, D.A., Tsuchida, C.A., Al-Shayeb, B., Banfield, J.F., Nogales, E., and Doudna, J.A. (2021). DNA interference states of the hypercompact CRISPR–CasΦ effector. Nature Structural & Molecular Biology 28, 652–661. 10.1038/s41594-021-00632-3.

53. Edgar, R.C. (2004). MUSCLE: multiple sequence alignment with high accuracy and high throughput. Nucleic Acids Research 32, 1792–1797. 10.1093/nar/gkh340.

54. Price, M.N., Dehal, P.S., and Arkin, A.P. (2010). FastTree 2 – approximately Maximum-Likelihood trees for large alignments. PLoS ONE 5, e9490. 10.1371/journal.pone.0009490.

55. Letunic, I., and Bork, P. (2024). Interactive Tree of Life (iTOL) v6: recent updates to the phylogenetic tree display and annotation tool. Nucleic Acids Research 52, W78–W82. 10.1093/nar/gkae268.

56. Wang, D., Zhang, F., and Gao, G. (2020). CRISPR-Based Therapeutic Genome Editing: strategies and in vivo delivery by AAV vectors. Cell 181, 136–150. 10.1016/j.cell.2020.03.023.

57. Khlebnikov, A., Keasling, J.D., Wanner, B.L., Skaug, T., and Datsenko, K.A. (2001). Homogeneous expression of the PBAD promoter in Escherichia coli by constitutive expression of the low-affinity high-capacity AraE transporter. Microbiology 147, 3241– 3247. 10.1099/00221287-147-12-3241.

58. Metcalf, W.W., Jiang, W., Daniels, L.L., Kim, S.-K., Haldimann, A., and Wanner, B.L. (1996). Conditionally replicative and conjugative plasmids CarryingLACZΑ for cloning, mutagenesis, and allele replacement in bacteria. Plasmid 35, 1–13. 10.1006/plas.1996.0001.

59. Strecker, J., Zhang, F., Ladha, A. CRISPR-associated transposase systems and methods of use thereof. International patent WO2020131862A1 (2020).

